# Reporters of TCR signaling identify arthritogenic T cells in murine and human autoimmune arthritis

**DOI:** 10.1101/474700

**Authors:** Judith F. Ashouri, Lih-Yun Hsu, Steven Yu, Dmitry Rychkov, Yiling Chen, Debra A. Cheng, Marina Sirota, Erik Hansen, Lisa Lattanza, Julie Zikherman, Arthur Weiss

**Author notes:** J.F.A and L.-Y.H. contributed equally to this work.

## Abstract

How pathogenic CD4 T cells in Rheumatoid Arthritis (RA) develop remains poorly understood. We used Nur77—a marker of T cell antigen receptor (TCR) signaling—to identify antigen-activated CD4 T cells in the SKG mouse model of autoimmune arthritis and in patients with RA. Using a fluorescent reporter of Nur77 expression in SKG mice, we found that higher levels of Nur77-eGFP in SKG CD4 T cells marked their autoreactivity, arthritogenic potential, and ability to more readily differentiate into IL-17 producing cells. The T cells with increased autoreactivity, nonetheless had diminished *ex vivo* inducible TCR signaling, perhaps reflective of adaptive inhibitory mechanisms induced by chronic auto-antigen exposure *in vivo*. The enhanced autoreactivity was associated with upregulation of IL-6 cytokine signaling machinery, which might in part be attributable to a reduced amount of expression of suppressor of cytokine signaling 3 (SOCS3)—a key negative regulator of IL-6 signaling. As a result, the more autoreactive GFP^hi^ CD4 T cells from SKGNur mice were hyper-responsive to IL-6 receptor signaling. Consistent with findings from SKGNur mice, *SOCS3* expression was similarly downregulated in RA synovium. This suggests that, despite impaired TCR signaling, autoreactive T cells exposed to chronic antigen stimulation exhibit heightened sensitivity to IL-6 which contributes to the arthritogenicity in SKG mice, and perhaps in patients with RA.

**Significance Statement:** How arthritis-causing T cells trigger rheumatoid arthritis (RA) is not understood since it is difficult to differentiate T cells activated by inflammation in arthritic joints from those activated through their TCR by self-antigens. We developed a model to identify and study antigen-specific T cell responses in arthritis. Nur77—a specific marker of TCR signaling—was used to identify antigen-activated CD4 T cells in the SKG arthritis model and patients with RA. Nur77 could distinguish highly arthritogenic and autoreactive T cells in SKG mice. The enhanced autoreactivity was associated with increased IL-6-receptor-signaling, likely contributing to their arthritogenicity. These data highlight a functional correlate between Nur77 expression, arthritogenic T cell populations, and heightened IL-6 sensitivity in SKG mice with translatable implications for human RA.

## Introduction

Rheumatoid arthritis (RA) is a chronic, destructive autoimmune disease that targets both joints and other organs. CD4 T cells have long been appreciated to play a crucial role in the pathogenesis of RA (1-4). Cellular and biochemical analyses of human CD4 T cells have revealed abnormal TCR signaling in RA patients (5-8). Surpisingly, CD4 T cells from patients with RA are hypo-responsive to TCR engagement *ex vivo* as evidenced by reduced calcium mobilization and IL-2 production (9, 10). This may be due, in part, to reduced TCRζ and LAT expression, and cellular changes associated with immune senescence (6, 7, 11-16). Yet, despite this, CD4 T cells in RA hyper-proliferate during clonal expansion and differentiate into effector cells that drive disease (17, 18). It is not currently understood how to reconcile these paradoxical observations of diminished TCR signaling in the setting of increased T cell proliferation and effector functions, nor is it clear whether this RA T cell phenotype is directly causal in disease pathogenesis or rather results from exposure to chronic inflammation. The inability to identify relevant arthritogenic T cells in patients and in murine disease models has limited our understanding of disease-initiating events in RA. In this report, we have developed a strategy to overcome this limitation by taking advantage of the dynamic expression pattern of Nur77(*Nr4a1*) in T cells.

*Nr4a1* is an immediate early gene that encodes the orphan nuclear hormone receptor Nur77. It is rapidly and robustly upregulated by TCR but not cytokine stimulation (19, 20). Therefore, Nur77 expression in murine and human T cells serves as a specific marker of TCR signaling, but is insensitive to cytokine stimulation (21-24). Detection of Nur77 expression can be used to identify T cells stimulated by direct self-antigen exposure prior to disease initiation as well as in the context of complex immune responses and chronic autoimmune diseases in which inflammatory cytokines can influence T cell phenotypes. Indeed, gene expression data show that endogenous *Nr4a1* transcripts are highly upregulated in autoantigen-specific CD4 T cells *in vivo* in the context of bona fide autoimmune disease (Immunological Genome Project; www.immgen.org, T.4.Pa.BDC)(25). This suggested to us that Nur77 expression could be employed to identify autoantigen-specific CD4 T cells in RA.

SKG mice harbor a profoundly hypomorphic variant of the Zap70 cytoplasmic tyrosine kinase, a molecule that is critical for proximal TCR signal transduction. As a consequence, SKG mice exhibit impaired thymocyte negative selection, giving rise to T cells with a more potentially autoreactive TCR repertoire (26). In response to either an innate immune stimulus in the form of zymosan injection or adoptive transfer of CD4 T cells into lymphopenic hosts, tolerance is broken, and mice develop an erosive inflammatory arthritis that resembles RA. For example, rheumatoid factor, anti-CCP antibodies and interstitial lung inflammation develop at the onset of disease (26). The SKG mouse provides a useful model to study early events in RA pathogenesis; not only does it capture many important features of human RA, but SKG also offers two advantages over other RA models. First, unlike the more commonly used collagen-induced arthritis (27) or K/BxN serum-transfer models of arthritis, which both bypass the initial break in TCR tolerance (28), the SKG mice exhibit a loss in central, and likely peripheral, tolerance that can be molecularly dissected. Hence this model uniquely allows us to study arthritis-causing T cells both before and during disease. Second, it recapitulates the paradoxical ability of RA CD4 T cells to differentiate into pathogenic effector cells despite impaired TCR signaling (6, 7, 11-18, 26).

A recent study of the SKG model has identified one specific arthritogenic TCR directed against a ubiquitous self-antigen (29), but it is not known whether rare antigen-specific T cell clones drive disease, or whether the entire pre-immune TCR repertoire has arthritogenic potential because it is highly autoreactive. Nor is it clear how tolerance of such clones is broken in the face of profoundly depressed TCR signaling in SKG mice. To address these questions we backcrossed the Nur77-eGFP BAC transgenic reporter line (in which eGFP protein expression is under the control of the *Nr4a1* regulatory region (24)) onto the SKG mouse model of arthritis. The resulting so-called SKGNur mice provided us with a tool to facilitate identification and study of arthritogenic CD4 T cells both before and after disease initiation. Despite impaired TCR signal transduction, the peripheral naïve CD4 T cells from SKG mice expressed higher levels of Nur77-eGFP relative to those of wild type mice. This suggested that even before disease initiation, SKG mice harbor a profoundly autoreactive T cell repertoire that exhibits increased downstream signaling despite impaired Zap70 function. We show here that the amount of Nur77-eGFP expression in CD4 T cells from SKG mice correlated with both their autoreactivity and their arthritogenicity. We found that the SKGNur CD4 T cells that expressed the highest level of the Nur77-eGFP reporter (GFP^hi^) differentiated more readily into IL-17 producing cells *in vivo*. While examining why GFP^hi^ cells were pre-disposed to differentiate into Th17 effector T cells, we observed profoundly enhanced IL-6-dependent signaling in GFP^hi^ CD4 T cells from SKG mice. Apparently, the autoimmune repertoire despite being coupled with impaired TCR signal transduction capacity in the SKG model resulted in reduced expression of suppressor of cytokine signaling 3 (SOCS3)—a key negative regulator of IL-6 signaling. Likewise, we found *SOCS3* expression was downregulated in RA synovium. This in turn suggests that, despite impaired TCR signaling, autoreactive T cells exposed to chronic antigen stimulation exhibit heightened sensitivity to IL-6 receptor signaling which contributes to their arthritogenicity in SKG mice, and perhaps in patients with RA.

## Results

### A fluorescent reporter of TCR signaling reveals and marks self-reactivity of the CD4 T cell repertoire in SKG mice

We sought to take advantage of a transcriptional reporter of *Nr4a1* expression, Nur77-eGFP BAC Tg, in order to examine antigen-specific signaling in pathogenic SKG T cells. Since Nur77 expression is induced downstream of TCR but not cytokine receptor signaling (21, 23, 24), it can serve as a specific reporter of antigen-receptor signaling in T cells *in vitro* or *in vivo* (21-24). We back-crossed the Nur77-eGFP BAC Tg reporter (24) into either wild type (WT) or SKG mice, both on the Balb/c background (hereafter referred to as WTNur or SKGNur, respectively). We confirmed that the introduction of the Nur77-eGFP BAC Tg reporter into the SKG mouse model did not alter their phenotype. Indeed, the SKGNur CD4 T cells continued to demonstrate dramatic hyporesponsiveness to *in vitro* polyclonal TCR stimulation, consistent with published data (26, 30), and developed a severe inflammatory and erosive arthritis after zymosan challenge (**Fig. S1A-C**).

It has previously been shown that thymic negative selection is quite impaired in SKG mice because TCR signaling is exceptionally dampened (26). As a result, highly autoreactive T cells, that would ordinarily be deleted, are instead positively selected. Although one SKG autoreactive TCR has been cloned (29), it remains unknown whether rare antigen-specific T cell clones drive disease, or whether the entire repertoire has arthritogenic potential because it is highly autoreactive. The Nur77-eGFP distribution in SKG mice is broad and overlaps with the distribution seen in normal mice (**Fig. 1A-B**). Following stimulation with zymosan, which induces arthritis, there is an enrichment of Nur77-eGFP expressing CD4 T cells with higher levels of GFP in draining popliteal nodes, but the distribution of GFP is still quite broad and largely overlaps with that of cells from untreated SKG mice. The Nur77-eGFP expression in CD4 T cells isolated from the arthritic joints of SKG mice, however, is substantially increased but still overlaps with the expression distribution in CD4 T cells from untreated mice (**Fig. 1A**, gating in **Fig. S1D**). This suggests that the T cell repertoire prior to and after zymosan stimulation has a broad distribution but the arthritogenic potential may not be limited to a narrow spectrum of autoreactive cells. This could account for the failure to date to identify a unique, dominant arthritogenic TCR in this model (29). We took advantage of SKGNur mice to study this further.

**Figure 1.**
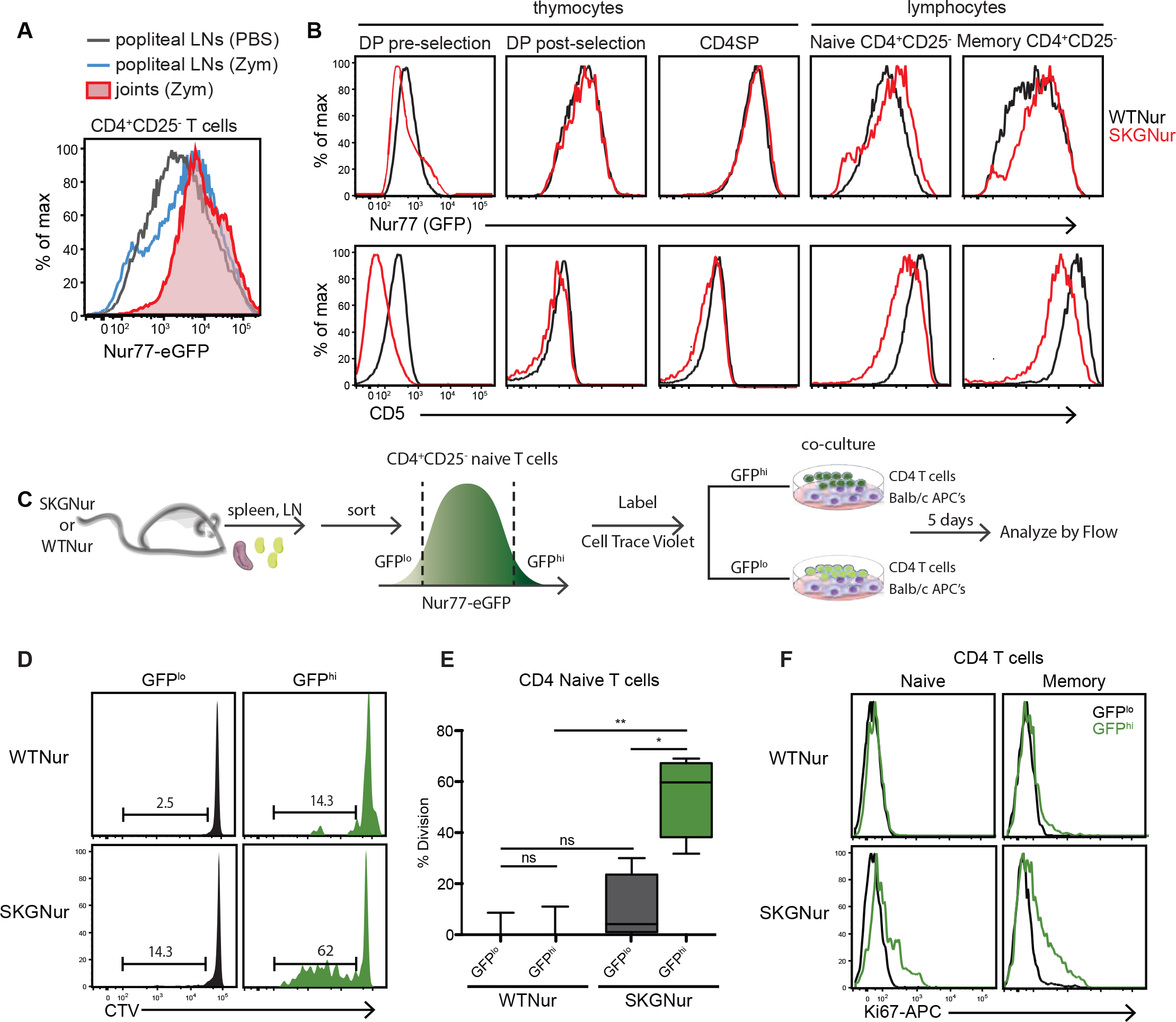
The Nur77 reporter of TCR signaling marks the selection of an autoreactive repertoire during thymic development with further pruning in the periphery. **(A)** Assessment of Nur77-eGFP fluorescence in SKGNur CD4^+^CD25^-^ T cells freshly isolated from popliteal LN ± PBS or zymosan and arthritic joints (+ zymosan). Data are representative of >10 mice in at least 4 independent experiments. **(B)** Comparison of expression of Nur77-eGFP and CD5 in pre-selection (CD5^int^TCRβ^int^) DP, CD4SPCD25^-^ thymocytes, and CD4^+^CD25^-^ naïve and memory peripheral T cells. Results are representative of 3 independent experiments with 6 mice in each group. **(C)** Depiction of AMLR workflow: **(D-E)** Sorted CD4^+^CD25^-^ naïve GFP^lo^ and GFP^hi^ T cells were loaded with cell trace violet dye (CTV) and co-cultured with APCs from Balb/c mice for 5 d. Cells were subsequently stained for T-cell markers and assessed for dye dilution by flow cytometry **(D)**. **(E)** Box plots represent % division of GFP^lo^ and GFP^hi^ naïve CD4 T cells as assessed using the Flow Jo v9 proliferation platform (center line, median; box limits, upper and lower quartiles; whiskers, 1.5x IQR). Representative histogram of **(D)** CTV dilution and **(E)** % cell divison from 4 experimental replicates (each containing 2-3 pooled mice). \**P* < 0.05, \*\**P* < 0.01, by Student’s t test. **(F)** Histograms represent Ki67 proliferation marker staining in GFP^lo^ and GFP^hi^ CD4^+^CD25^-^ naïve (CD62L^hi^CD44^lo^) or memory (CD44^hi^CD62L^lo^) T cells from WTNur or SKGNur mice.

We first used the Nur77-eGFP reporter to assess differences in TCR signaling between WT and SKG T cells during T cell development, as it seemed likely that a more autoreactive repertoire was being positively selected by self pMHC in SKG mice. The surface marker CD5 is dependent upon p56^lck^ activity throughout T cell development and is frequently used as an indicator of TCR signaling strength during and after thymic development (31). We assessed both CD5 and Nur77-eGFP expression at sequential stages of T cell development in WTNur and SKGNur mice. Consistent with impaired TCR signaling in SKG mice as identified in previous studies, pre-selection double-positive (DP) thymocytes from SKGNur mice exhibited lower levels of both GFP and CD5 relative to those of WTNur (**Fig. 1B** left panel). Prior work has identified a minimal threshold of TCR signaling that is necessary for positive selection (32). Consistent with this, after the positive selection checkpoint, GFP and CD5 were markedly upregulated in post-selection thymocytes in both WTNur and SKGNur mice. Of note, GFP expression levels in SKGNur CD4SP thymocytes became equivalent to WTNur, whereas CD5 expression remained subtly reduced in SKGNur CD4SP thymocytes (**Fig. 1B**). Since TCR signaling is markedly dampened in SKG mice, high GFP expression in SKGNur CD4SP thymocytes implies selection of a highly autoreactive repertoire where strong stimulation by self-peptide MHC (self-pMHC) complexes compensates for impaired signal transduction in order to support positive selection (32).

Surprisingly, we found that peripheral naïve and memory CD4 lymphocytes from SKGNur mice had somewhat higher GFP levels than WTNur controls (**Fig. 1B**, two right-hand panels). Note that Tregs, which are selected on the basis of self-reactivity during development and consequently express high levels of Nur77 (23), were excluded in this analysis. This suggested to us that autoreactive SKG T cells were either exposed to chronic auto-antigen stimulation in the periphery or underwent further repertoire pruning under the influence of auto-antigens and weak TCR signaling, or both. In contrast, CD5 surface expression was lower in SKGNur peripheral CD4 T cells compared to controls (**Fig. 1B**). It is unclear why CD5 and Nur77 expression levels were discordant following positive selection and in the periphery.

However, CD5 is frequently used as a marker of proximal TCR signaling strength involving interactions with Lck and Cbl with the CD5 cytoplasmic tail, (9, 33, 34). In contrast, Nur77 is reflective of transcriptional activation which is regulated by tonic and integrated downstream TCR signaling pathways (35). Thus, we used Nur77-eGFP as a marker of TCR signaling strength in SKG peripheral T cells as it appeared to be a reliable marker of, not only strength of transduced TCR signal, but integrated TCR signaling reflective of antigen encounter *in vivo* over time.

### Nur77-eGFP high expressing SKGNur CD4 T cells demonstrate increased autoreactivity

In view of the discordant expression of Nur77-eGFP and CD5 in peripheral CD4 T cells from SKGNur mice and the broad distribution of Nur77-eGFP expression, we sought to take an independent approach to probe the putative self-reactivity of the mature repertoire and whether Nur77-eGFP expression marked differences in the self-reactivity of this repertoire. We compared the response of CD4^+^CD25^-^ naïve T cells sorted for the 10% highest and 10% lowest Nur77-eGFP expression (hereafter referred to as GFP^hi^ and GFP^lo^ respectively) to determine whether reporter expression reflected repertoire differences and/or interactions with self-pMHC among the polyclonal CD4 T cells. Sorted CD4^+^CD25^-^ GFP^hi^ and GFP^lo^ naïve T cells from SKGNur and WTNur mice were challenged in an *in vitro* autologous mixed lymphocyte reaction (AMLR). Sorted cells were stained with cell trace violet (CTV) and incubated *in* vitro with WT Balb/c splenocytes for 5 days in culture and proliferation was assessed by CTV dye dilution (**Fig. 1C**). WTNur T cells are not expected to mount a response in this assay because of efficient tolerance induction during thymic negative selection. Indeed, as expected, neither GFP^hi^ nor GFP^lo^ CD4 T cells from WTNur mice mounted substantial proliferative responses. However, SKGNur GFP^hi^ CD4^+^CD25^-^ naive T cells exhibited at least a four-fold greater *in vitro* proliferative response to autologous Balb/c APCs compared to SKG-GFP^lo^ CD4 T cells (**Fig. 1D-E**). This ability to proliferate in an autologous AMLR suggests that SKGNur GFP^hi^ CD4 T cells indeed are enriched for cells with self-reactive TCRs, and that the degree of autoreactivity is sufficient to overcome impaired TCR signal transduction conferred by the SKG *Zap70* hypomorphic allele.

We then assessed GFP^hi^ and GFP^lo^ CD4 T cell populations from WTNur and SKGNur mice for evidence of an *in vivo* correlate of the *in vitro* proliferative response. As shown in **Fig. 1F**, we compared Ki-67 expression in both naïve and memory CD4 T cells from WTNur and SKGNur mice. As expected, under steady-state conditions, virtually no naïve WT T cells were proliferating, and Ki67 expression was virtually undetectable in both GFP^hi^ and GFP^lo^ CD4 T cells from WTNur mice. However, a substantial proportion of naïve and memory T cells from SKG mice were Ki67+, consistent with a previous BrdU-labeling study in which SKG CD4 T cells were shown to have high *in vivo* proliferative activity (36). Importantly, the proliferating fraction was almost exclusively found among GFP^hi^ cells, suggesting that Nur77-eGFP expression indeed marks a functionally distinct CD4 T cell population in the periphery of SKGNur mice, presumably driven at least in part by stronger recognition of self-pMHC *in vivo* and *in vitro*. Moreover, this functional heterogeneity is identifiable by the extent of the Nur77-eGFP reporter expression before frank disease onset.

### Nur77-eGFP expression in SKGNur CD4 T cells strongly correlates with their arthritogenic potential

Since the Nur77-eGFP reporter appeared to mark more autoreactive T cells, we next asked whether this, in turn, correlated with their capacity to drive arthritis. We used an adoptive transfer model of arthritis (26, 37, 38), to determine whether reporter expression identified a more arthritogenic population of CD4 T cells. In this model, CD4 T cells transferred from SKG mice, but not sera or CD8 or B cells, are sufficient to cause arthritis (26). As delineated in **Fig. 2A**, CD4^+^CD25^-^ GFP^hi^ and GFP^lo^ naïve T cells were sorted and adoptively transferred into immunodeficient SCID recipients. Initially we found that, with sufficient time (>6 weeks post transfer), both GFP^hi^ and GFP^lo^ CD4 SKG T cells were capable of producing arthritis upon transfer to lymphopenic hosts, while WT T cells were not. This suggested that the entire SKG T cell repertoire harbors T cells with pathogenic potential. Therefore, we next sought to evaluate whether a difference in onset and severity could be histologically detected. Indeed, transfer of CD4^+^CD25^-^ naïve T cells expressing the highest levels of GFP (GFP^hi^) resulted in an earlier onset (14 days after adoptive transfer) and more severe arthritis in recipient SCID mice than did the transfer of GFP^lo^ naïve CD4 T cells (**Fig. 2B-C**). This was characterized by increased synovial proliferation, inflammatory infiltrates, and early cartilage destruction (white arrows). This was not observed in SCID mice that received either SKGNur GFP^lo^ CD4^+^CD25^-^ naïve T cells, or WTNur GFP^hi^ or GFP^lo^ CD4^+^CD25^-^ naïve T cells, where the thin layer of synovial lining is maintained and infiltrating inflammatory cells are absent (**Fig. 2B-C**, black arrows). To assess whether these results were unique to the naïve SKGNur GFP^hi^ CD4 T cell population, the adoptive transfer experiment was repeated after sorting for GFP^hi^ and GFP^lo^ CD4^+^CD25^-^ memory T cells (**Fig. S2A**). We likewise found that the SKGNur GFP^hi^ CD4^+^CD25^-^ memory T cells were capable of producing a more severe and earlier onset of arthritis than SKGNur GFP^lo^ T cells or Balb/c WTNur control T cells (**Fig. S2B**). We conclude that arthritogenicity is a property of the entire SKG CD4 T cell repertoire, but that T cells have varying arthritogenic potential, and most importantly, that this arthritogenic potential can be identified by the amount of Nur77-eGFP expression and reflects their degree of self-reactivity.

**Figure 2.**
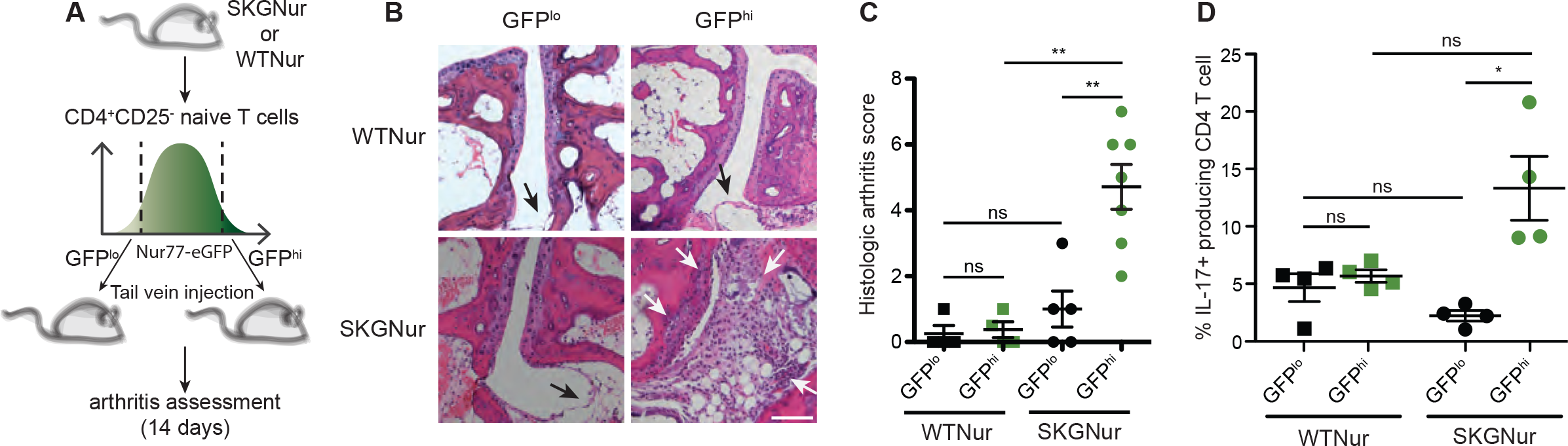
Nur77-eGFP marks arthritogenicity of CD4 T cells in SKGNur mice. **(A)** The 5% highest and 5% lowest Nur77-eGFP purified CD4 T cells sorted on naïve markers (CD62L^hi^CD44^lo^) and depleted of Treg marker CD25 were adoptively transferred into SCID recipients. **(B)** Joints were assessed by histologic staining 14 d following adoptive transfer and scored in a blinded fashion. Blacks arrows indicate normal synovial lining and white arrows indicate synovial proliferation and joint destruction, white scale bar, 100μ. **(C)** Dot plot of the histologic score from SCID recipients that received GFP^lo^ (black) or GFP^h^ (47) CD4 naïve T cells from WTNur (squares) or SKGNur (circles) mice. Each dot represents one biologically independent sample pooled from 3 experiments, mean ± SEM. **(D)** Dots represent % IL-17 producing GFP^+^ CD4 T cells from SCID recipient splenocytes stimulated *in vitro* with PMA and ionomycin for 5 h, mean ± SEM.

Since the GFP^hi^ population in SKGNur mice had greater arthritic potential, we asked whether the GFP^hi^ population exhibited distinct T cell effector functions from those of the GFP^lo^ population in SKGNur mice. To address this question, we determined their ability to generate IL-17 *in vitro* by stimulating sorted primary naïve T cell populations with plate-bound anti-CD3 and CD28. Notably, we found that the percentages of IL-17 producing cells were 5-to 10-fold higher in the SKGNur GFP^hi^ population than in the SKGNur GFP^lo^ population. However, the difference in IL-17 production between GFP^hi^ and GFP^lo^ populations from WTNur mice was almost indistinguishable (**Fig. S2C**). To determine whether this difference could be detected in our adoptive transfer model, transferred naïve or memory CD4 T cells recovered from the recipient SCID mice were stimulated for 5 hours *ex vivo* with PMA and ionomycin in the presence of brefeldin A to identify Th17 effector T cells capable of secreting IL-17. CD4 T cells recovered from SCID recipient mice that received SKGNur GFP^hi^ naïve or memory CD4 T cells had a significantly higher percentage of such IL-17-producing T cells than their GFP^lo^ counterparts (**Fig. 2D, Fig. S2D**). This increase in IL-17 producing cells was not observed in the CD4 T cells recovered from SCID mice that received WTNur GFP^hi^ or GFP^lo^ CD4 T cells. Collectively, our results suggest that the arthritogenic capacity of SKG CD4 T cells can be distinguished by their relative expression of Nur77-eGFP before disease onset, and corresponds to increased autoreactivity and proclivity to differentiate into pathogenic IL-17-secreting effectors in vitro and in vivo.

### Autoreactive Nur77-eGFP^hi^ SKG T cells signal poorly in response to TCR stimulation

We next sought to understand why arthritogenic GFP^hi^ SKG T cells preferentially differentiated into Th17 effectors. Since GFP levels reflect integrated TCR signaling, we hypothesized that differences in TCR signaling by this population of T cells may contribute to their Nur77-GFP amounts (**Fig. 3A**). We already showed that GFP^hi^ SKG T cells are more self-reactive and therefore encounter more antigen *in vivo* (**Fig. 1C-D**). We next wanted to determine whether cell-intrinsic capacity to signal downstream of their TCRs differed between GFP^hi^ and GFP^lo^ T cells. To test this, we probed signaling events downstream of the TCR. By removing cells from *in vivo* antigen exposure and treating them with the polyclonal stimulus α-CD3 *in vitro*, we were able to bypass any differences in TCR repertoire between the GFP^hi^ and GFP^lo^ CD4 T cell subsets. We first analyzed TCR-induced increases in cytoplasmic free calcium in peripheral naïve T cells. SKGNur CD4^+^CD25^-^ T cells irrespective of GFP expression demonstrated profoundly dampened TCR-induced increases in cytoplasmic free calcium, consistent with the hypomorphic SKG Zap70 allele (**Fig. 3B, Fig. S1**). The calcium responses of GFP^hi^ SKGNur CD4 T cell responses were even more dampened than those of GFP^lo^ CD4 T cells (**Fig. 3A**). Interestingly, an analogous trend was seen with GFP^hi^ WTNur CD4 T cells relative to GFP^lo^ WTNur CD4 T cells, suggesting that increased integrated TCR signaling input is associated with dampened downstream TCR signal transduction.

**Figure 3.**
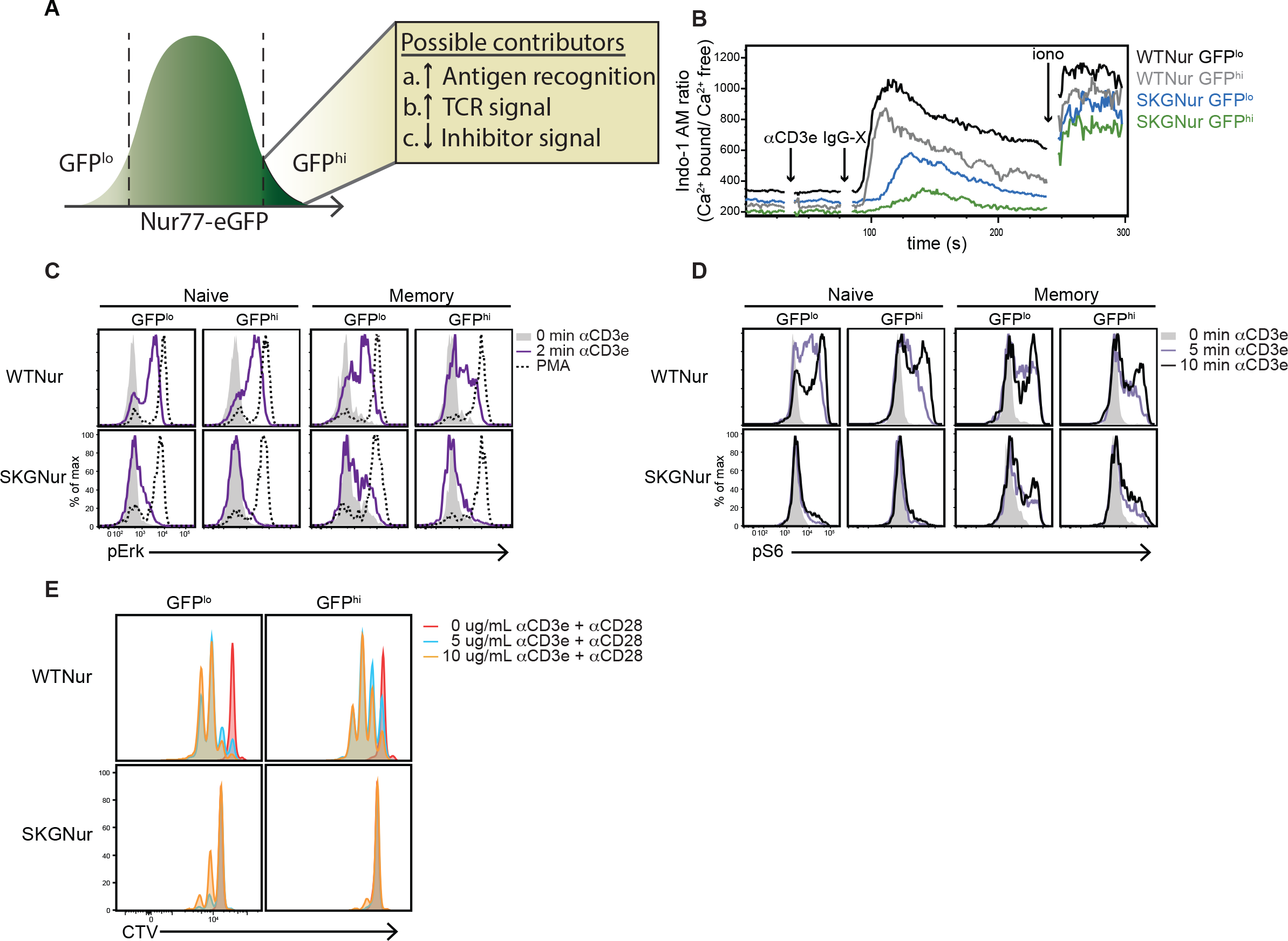
Autoreactive Nur77-eGFP^hi^ SKG T cells signal poorly. **(A)** Integrated TCR signaling inputs reflect Nur77-eGFP levels. **(B)** TCR-induced calcium increases in GFP^lo^ and GFP^hi^ negatively selected CD4^+^CD25^-^ T cells from WTNur and SKGNur mice. Data are representative of 4-5 mice in each group from 3 independent experiments. **(C-D)** Histograms represent p-Erk **(C)** and p-S6 **(D)** levels analyzed in GFP^lo^ and GFP^hi^ CD4^+^CD25^-^ T cells from WTNur and SKGNur mice by flow cytometry after stimulation with α-CD3ε. Data are representative of at least 8 mice in each group from 4 independent experiments. **(E)** Sorted CD4^+^CD25^-^ naïve GFP^lo^ and GFP^hi^ T cells were loaded with CTV and stimulated with plate bound α-CD3ε and 2 ug/mL αCD28 for 3 d. Cells were subsequently assessed for dye dilution by flow cytometry. Data are representative of 2 pooled mice in each group from 2 independent experiments.

Analysis of TCR-induced Erk phosphorylation in purified peripheral CD4 T cells similarly revealed that both GFP^hi^ and GFP^lo^ CD4^+^CD25^-^ SKGNur T cells had a severe impairment in Erk phosphorylation after polyclonal TCR stimulation. This was observed in both the memory and naïve peripheral CD4 T cell compartments (**Fig. 3C, Fig. S1**). By contrast, responses to PMA which bypasses TCR signaling to Erk activation were normal, consistent with the well-defined role of Zap70 in regulating only proximal TCR signal transduction. In WTNur mice, GFP^hi^ CD4 T cells appeared to have subtly decreased signaling capacity compared to GFP^lo^ T cells, but the responses were still largely preserved, contrasting with the markedly impaired SKG responses (**Fig. 3C**). Similarly, GFP^hi^ more than GFP^lo^ CD4^+^CD25^-^ SKGNur T cells had severely dampened PI3K/Akt signaling downstream of polyclonal TCR stimulation as analyzed by S6 phosphorylation at Ser240/244 (**Fig. 3D**). Likewise, though WTNur GFP^hi^ CD4 T cells exhibited a slightly dampened response compared to GFP^lo^ CD4 T cells, PI3K/Akt signaling downstream of TCR stimulation was much more robust in WTNur cells when compared to SKGNur cells (**Fig. 3D**). Consistent with our calcium mobilization and phosphoflow studies, sorted GFP^lo^ CD4^+^CD25^-^ naïve CD4 T cells from both SKGNur and WTNur mice proliferated more robustly to platebound αCD3 stimulation than GFP^hi^ naïve CD4 T cells (**Fig. 3E**). Moreover, SKGNur CD4 T cells exhibited a clear proliferative defect in response to platebound αCD3 stimulation as previously described (30), which stands in marked contrast to SKGNur GFP^hi^ naïve CD4 T cells’ ability to proliferate in response to self-antigen (**Fig. 1D-E**). Therefore, these results indicate that differences in GFP expression between the most and least arthritogenic SKGNur CD4 T cells is not attributable to signal transduction machinery, but rather reflects TCR autoreactivity to self-pMHC (antigen recognition). It is possible that downmodulation of TCR signal strength in GFP^hi^ relative to GFP^lo^ CD4 T cells could represent a compensatory peripheral tolerance mechanism to prevent inappropriate activation of self-reactive clones. Collectively, these results suggest that differences in the measured TCR signal transduction events do not account for unique propensity of GFP^hi^ CD4 T cells from SKGNur mice to differentiate *in vivo* and *in vitro* into Th17 effectors.

### Upregulation of several inhibitory receptors in SKG GFP^hi^ cells

It was quite surprising that GFP^hi^ CD4 T cells from SKGNur mice were able to exert their effector functions leading to joint immunopathology despite their profound impairment to *in vitro* TCR signal transduction (**Fig. 3**). To account for the discrepancy between arthritogenic and signaling potential of the GFP^hi^ cells as detected here, we next asked whether GFP^hi^ CD4 T cells from SKGNur mice expressed much lower levels of inhibitory receptors, thereby permitting them to expand more rapidly in a lymphopenic environment, or whether they expressed higher levels of inhibitory receptors, further compromising their impaired TCR signaling capacity compared to GFP^lo^ CD4 T cells. To address this, we examined the surface expression of several inhibitory receptors (PD-1, TIGIT, CD73) which have been shown to dampen downstream TCR signaling, the latter of which marks ‘anergic’ T cell populations (39). GFP^hi^ CD4 T cells expressed higher levels of these inhibitory receptors than their GFP^lo^ counter parts, again suggesting that this may represent a compensatory tolerance mechanism to constrain self-reactive T cells (**Fig. 4**). However, higher expression levels of inhibitory receptors observed in the GFP^hi^ population were not unique to SKGNur T cells since WTNur GFP^hi^ CD4 T cells also upregulated inhibitory receptors compared to WTNur GFP^lo^ CD4 T cells. This is in line with a recent report demonstrating that *NR4A* gene expression positively correlated with *PDCD1* (PD-1), TIGIT, and other inhibitory receptors in the setting of chronic antigen stimulation as part of a T cell-intrinsic program of hyporesponsiveness (40). This may in part account for the relative hyporesponsiveness we observed in GFP^hi^ T cells (**Fig. 3**). Therefore it is all the more striking that SKGNur GFP^hi^ CD4 T cells upregulate inhibitory pathway receptors and yet, despite this, exhibit the capability to proliferate in response to autologous APCs, differentiate into IL-17 producing cells, and cause more severe arthritis.

**Figure 4.**
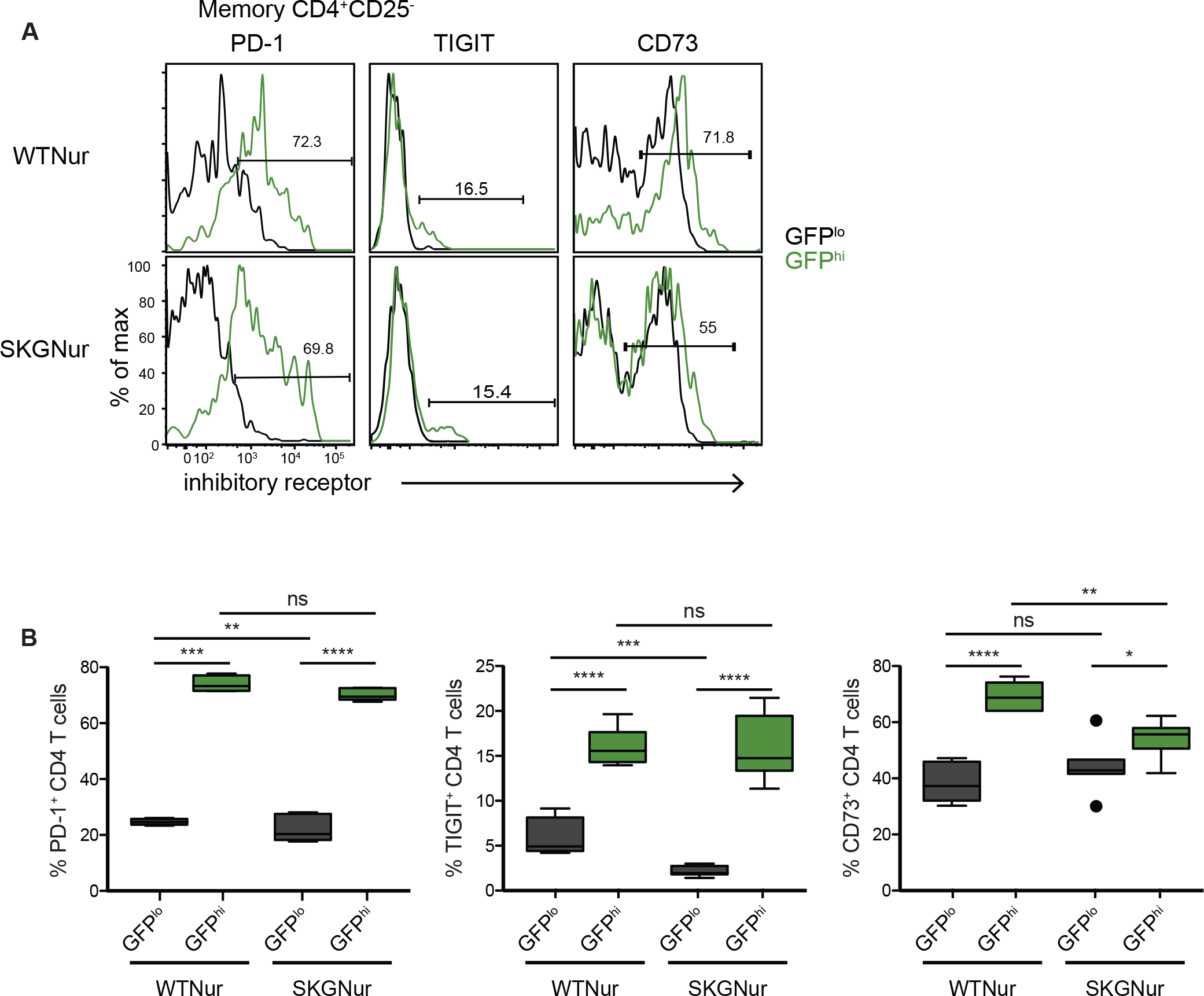
Inhibitory receptors are upregulated in GFP^hi^ CD4 T cells. **(A)**Expression levels of inhibitory receptors assessed by flow cytometry in memory CD4^+^CD25^-^ cells of GFP^lo^ and GFP^hi^ populations from WTNur or SKGNur mice. Results shown are representative of 2 independent experiments. (B) Quantification of % positive inhibitory receptor levels. Box plots describe the median and inter-quartile range (IQR), whiskers extend to the largest value, but no further than 1.5*IQR, data points beyond whiskers are outlying points. ns: *P* > 0.05, \**P* < 0.05, \*\**P* < 0.01, \*\*\**P* < 0.001, \*\*\*\**P* < 0.0001, by two-tailed Student’s t test.

### Enhanced IL-6 response in GFP^hi^ CD4 T cells from SKGNur mice

Th17 differentiation depends not only on TCR signaling in response to pMHC, but also upon polarizing cytokines. Given the critical role of IL-6 in the development of arthritogenic Th17 cells in SKG mice (36), the differential responsiveness to IL-6 signaling might, at least in part, account for differences in the arthritogenic and effector potential of GFP^hi^ and GFP^lo^ SKGNur CD4 T cells. We examined *in vitro* responsiveness of SKG and WT CD4 T cells to IL-6. Phospho-flow cytometric analyses revealed that SKGNur GFP^hi^ CD4 T cells have higher basal levels of phosphorylated-STAT3 than GFP^lo^ CD4 T cells (**Fig. 5A-B**). This difference was not observed amongst comparable populations from unstimulated WTNur CD4 T cells. Interestingly, inducible STAT3 phosphorylation in response to IL-6 stimulation in the GFP^hi^ populations was more robust than that of the GFP^lo^ populations but this was observed in both SKGNur and WTNur mice. The overall response to IL-6 stimulation in both the SKGNur GFP^hi^ and GFP^lo^ CD4 T cells was much greater than in the comparable populations obtained from WTNur CD4 T cells (**Fig. 5A-B**). To investigate whether IL-6 hyper-responsiveness in SKG mice was due to differences in total STAT3 protein expression levels, we first compared the expression levels of STAT3 between WT and SKG mice. As shown in **Fig. 5C**, we found that STAT3 expression was comparable between WT and SKG CD4 T cells.

**Figure 5.**
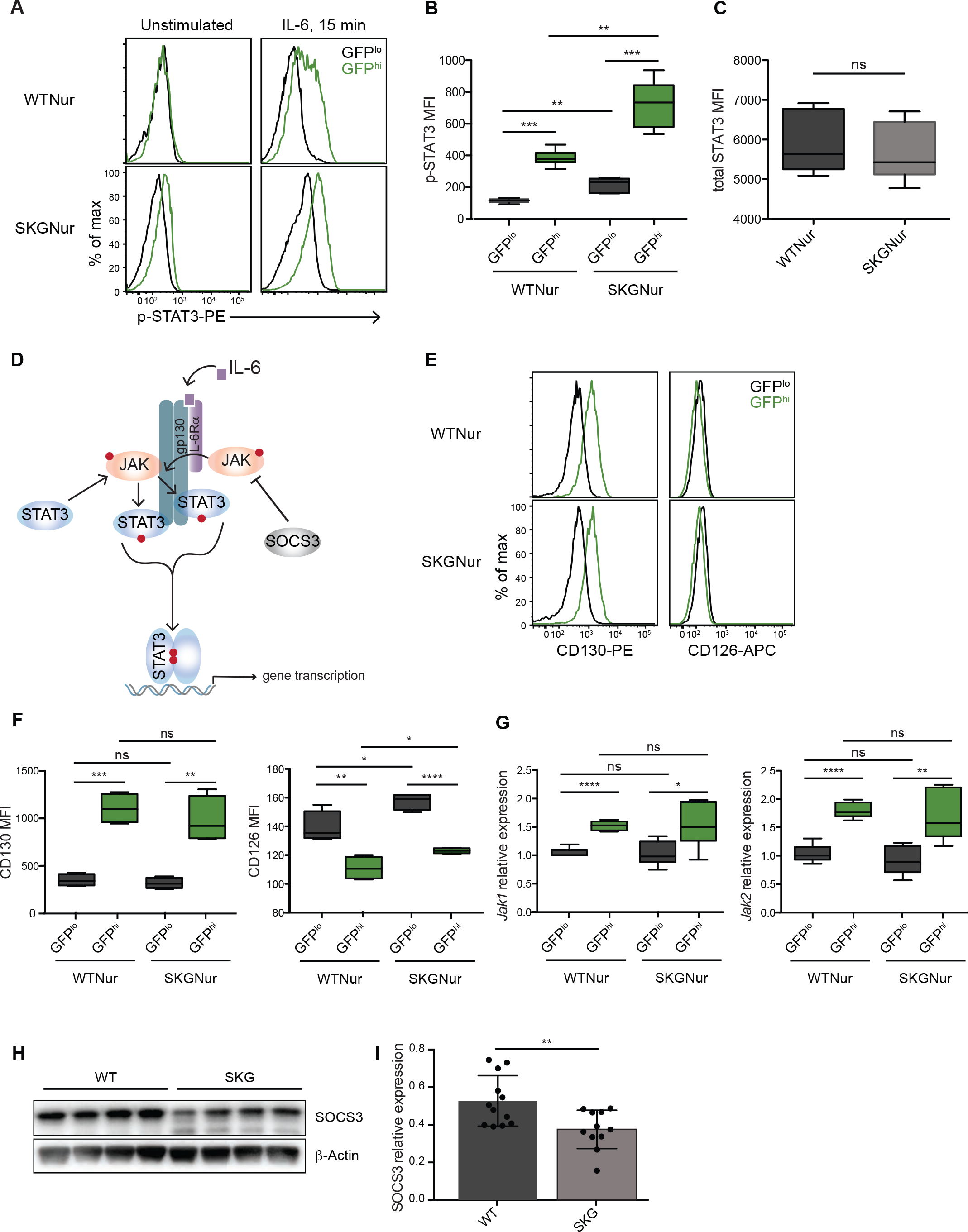
Enhanced IL-6 response in GFP^hi^ CD4 T cells from SKGNur mice. **(A)** Levels of p-STAT3 in GFP^lo^ and GFP^hi^ CD4 T cell populations from WTNur and SKGNur mice ± IL-6 treatment for 15 min. **(B)** Quantification of p-STAT3 mean fluorescence intensity (MFI) upon IL-6 stimulation from **(A)**. Data are pooled from 2 independent experiments, n=6 mice per group. **(C)** Quantification of total STAT3 MFI in WTNur (n=5) and SKGNur (n=10) mice. **(D)** IL-6 signaling pathway **(E)** Representative FACS analysis of CD130 and CD126 surface expression. **(F)** Quantification of CD130 and CD126 MFI. n=4 per group from 2 independent experiments. **(G)** Real-time RT-PCR measuring *Jak1* and *Jak2* mRNA levels in sorted GFP^lo^ and GFP^hi^ CD4 T cells from WTNur and SKGNur mice. n=5-7 per group from 2 independent sorts. All box plots represent quantification of MFI or RNA relative expression (center line, median; box limits, upper and lower quartiles; whiskers, 1.5x IQR). **(H)** Immunoblot analysis of SOCS3 and β-actin in unstimulated negatively selected CD4 T cells from SKG and WT mice following lysis with NP-40. Data are representative of three experiments. **(I)** Quantification of SOCS3 protein levels normalized to β-actin pooled from three immunoblot experiments as shown in **(H)** with 11-12 mice in each group. Ns: *P* > 0.05, \**P* < 0.05, \*\**P* < 0.01, \*\*\**P* < 0.001, \*\*\*\**P* < 0.0001, by two-tailed Student’s t test.

We next sought to examine the IL-6 pathway in greater detail (**Fig. 5D**). We first analyzed the expression of both components of the IL-6 receptor, the ligand-binding IL-6Rα chain CD126 and the IL-6 family common receptor subunit gp130 (CD130) which is essential for signal transduction following cytokine engagement (41). As shown in **Fig. 5E-F**, the GFP^hi^ populations expressed significantly higher amounts of CD130 than did the GFP^lo^ populations. However, there was no significant difference in CD130 expression between WTNur and SKGNur mice. We did observe minimal differential expression of CD126 protein between the GFP^hi^ and GFP^lo^ populations (**Fig. 5E-F**), although the impact of the very small differences is not clear. These results indicate that the GFP^hi^ and GFP^lo^ populations differ in their expression of CD130, perhaps contributing to relatively increased IL-6 responses by GFP^hi^ T cells.

Since the differences in the receptor subunits did not seem to account for the greater IL-6 response in SKGNur mice compared to WTNur controls, we further assessed key signaling molecules downstream of the IL-6 signaling pathway such as JAK1, JAK2, and SOCS3. Similar to surface expression of CD130, GFP^hi^ CD4 T cells expressed higher amounts of JAK1 and JAK2 mRNA than did GFP^lo^ CD4 T cells (**Fig. 5G**). Nevertheless, the expression levels of JAK1 and JAK2 transcripts were comparable between the respective CD4 T cell populations isolated from GFP populations in WTNur and SKGNur mice (**Fig. 5G**). Thus, increased expression of CD130, JAK1 and JAK2 in GFP^hi^ CD4 T cells might contribute to enhanced IL-6 response in this population, but both WTNur and SKGNur shared these properties. Immunoblot analyses indicated that SOCS3 protein, a key negative regulator of IL-6 signaling (42), was downregulated in SKGNur CD4 T cells (**Fig. 5H-I**). Our results suggest that the hyper-responsiveness to IL-6 observed in autoreactive SKGNur CD4 T cells exposed to chronic antigen stimulation may be due at least in part to decreased SOCS3-dependent suppression. We propose that SKGNur GFP^hi^ CD4 T cells exhibit uniquely robust IL-6 responses because they both upregulate the IL-6 proximal signaling machinery and coordinately express lower levels of the negative regulator SOCS3. Other regulators of this pathway could also influence the differential IL-6R responses of T cells from the WT vs SKG genotypes.

### Nur77 identifies increased frequencies of antigen-stimulated CD4 T cells in arthritic SKG synovium

A central challenge in isolating bona-fide antigen-activated T cells from inflamed arthritic joints from patients with rheumatoid arthritis is differentiating these from T cells activated via cytokines in the inflammatory milieu (43). Recently, we developed technologies to detect induction of endogenous Nur77 protein following TCR and BCR stimulation in mouse and human T cells. Since Nur77 is induced downstream of TCR but not cytokine receptor signaling (**Fig. 6A**) (21, 23, 24), and we observed increased expression of Nur77-eGFP in arthritic joints of SKG mice (**Fig. 1A**), we determined whether endogenous Nur77 protein was also upregulated in T cells after arthritis induction by zymosan challenge in SKG mice. Indeed, when examining the CD4 T cell population infiltrating arthritic SKG joints after zymosan treatment, we found upregulation of endogenous Nur77 protein in a subset of CD4 T cells from SKG arthritic joints compared to non-arthritic PBS treated SKG mice (**Fig. 6B,** gating in **Fig. S1D**). Consistent with intracellular Nur77 expression, Nur77 transcript levels (*Nr4a1*) were increased approximately 6-fold in the population of joint infiltrating CD4 T cells from zymosan-treated SKG mice (**Fig. 6C**).

**Figure 6.**
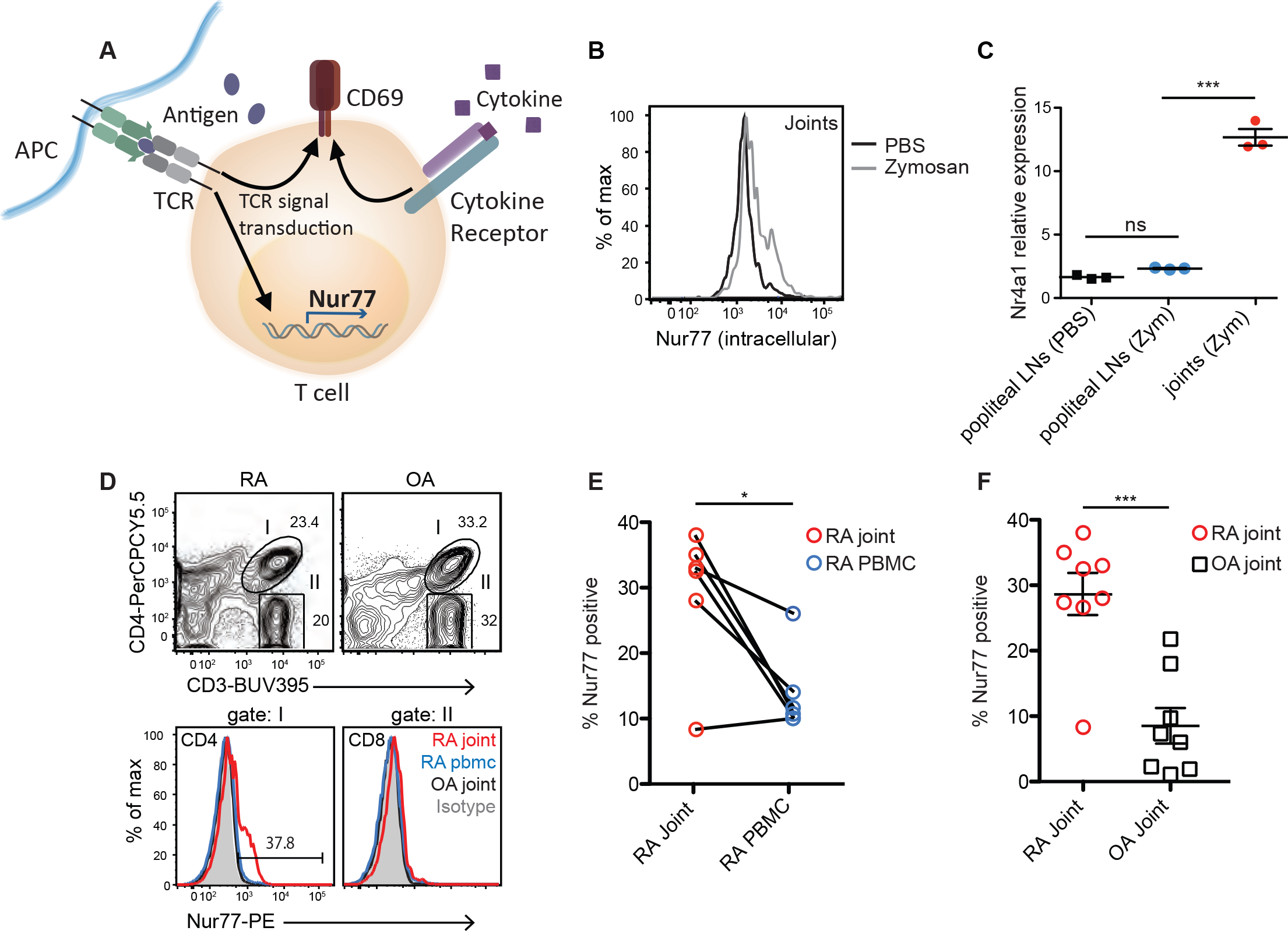
A reporter of TCR signaling can be used to identify antigen-activated CD4 T cells in human RA synovium. **(A)** Depiction of Nur77 induction by TCR signal transduction, but not cytokine signaling. **(B)** Histogram shows the overlay of endogenous Nur77 protein levels in the infiltrating CD4^+^CD25^-^ T cells from joints of SKGNur mice treated with PBS or zymosan. Data are representative of >10 mice in at least 5 independent experiments. **(C)** Comparison of mRNA transcript levels of Nr4A1 (Nur77) in sorted CD4^+^CD25^-^ T cells from LNs or joints of SKGNur mice treated with either PBS or zymosan. Each dot represents a biologically independent sample pooled from 2 independent experiments with 3 mice per group, mean ± SEM, Repeated measures ANOVA,\*\*\**P* ≤ 0.001. **(D)** Top panels, flow cytometry plots of RA and OA synovial tissue T cells demonstrate T cell populations from LiveCD45+ RA & OA synovium. Bottom histograms represent Nur77 expression in CD4 (gate I) & CD8 (gate II) T cells from synovial RA (red line) and OA (green line) tissue, RA PBMC’s (blue line), and isotype control (filled in grey histogram). **(E-F)** Dot plots demonstrate % Nur77 positive CD4 T cells (gate I) from paired RA PBMCs and joints, n=6, \**P* ≤ 0.05, by paired two-tailed-Student’s t test **(E)** and % Nur77 positive CD4 T cells from 8 RA and 8 OA synovial tissue, \*\*\**P* ≤ 0.001, by two-tailed-Student’s t test **(F)**. Each dot represents a biologically independent sample, mean ± SEM.

The discrepancy between greater percentage of CD4 T cells expressing high GFP amounts (**Fig. 1A**) compared to endogenous Nur77 may be due to the long *in vivo* half-life of GFP in comparison to endogenous Nur77 (22), but does also suggest a high degree of heterogeneity in the autoreactivity of the CD4 T cell population within the arthritic joint. Together, these data show that zymosan challenge in SKG(Nur) mice, which triggers activation of autoreactive CD4 T cells, led to upregulation of a population of cells that expressed higher levels of endogenous Nur77 expression in the inflamed joints. Since Nur77 marks antigen-dependent TCR signaling specifically, and the half-life of endogenous Nur77 is short (44), these results suggest that the joint-infiltrating CD4 T cells in SKG mice are recognizing an intra-articular antigen(s).

### Nur77 protein, a specific indicator of TCR signaling, identifies increased frequencies of antigen-stimulated CD4 T cells in human RA synovium

Utilizing methodology we optimized for intracellular Nur77 staining in human lymphocytes (21), we used endogenous Nur77 protein expression to determine whether joint-infiltrating CD4 T cells in human RA patients might be recognizing an intra-articular antigen, and as proof-of-concept to establish whether this marker could in turn be subsequently used to identify arthritogenic TCR specificities in RA patients. We analyzed Nur77 expression in synovial T cells from eight patients undergoing a surgical joint procedure that met the American College of Rheumatology classification criteria for seropositive (rheumatoid factor or anti-CCP antibody positive) RA (**Table S1**). We found a significant increase in a fraction of Nur77-positive CD4 T cells isolated from synovial tissue from patients with seropositive RA compared to matched PBMCs from the same donor (**Fig. 6D** and **6E,** human gating in **Fig. S3A**), suggesting an enrichment of antigen-reactive cells in the joint. The expression of Nur77 protein in CD4 T cells obtained from the joints of RA patients far exceeded those isolated from synovial tissue of patients with a diagnosis of osteoarthritis (OA), a non-autoimmune degenerative form of arthritis (**Fig. 6D, 6F, Fig. S3A**). Importantly, the enrichment of endogenous Nur77 was only observed in synovial infiltrating CD4 T cells and not in CD8 T cells in RA synovium (**Fig. 6D**), consistent with the genetic link to specific MHC Class II alleles and the role of CD4 T cells in the pathogenesis of RA (3-6, 8). These results support the hypothesis that pathogenic CD4 T cells are enriched in RA joints and are likely recognizing an intra-articular auto-antigen(s).

### SOCS3 transcripts are downregulated in RA synovial tissue

Similar to the SKG model, the pleiotropic cytokine IL-6 is thought to contribute to the differentiation of Th17 cells in human RA, and targeting the IL-6 receptor with clinically used humanized monoclonal antibodies leads to RA disease improvement (45, 46). Because SOCS3 induction is critical to prevent prolonged and enhanced IL-6 signaling *in vivo* (47), we wanted to investigate whether SOCS3 levels may be inappropriately reduced in patients with RA, creating a permissive environment for IL-6 signaling as seen in the SKG mouse model of arthritis. Indeed, Ye *et al.* found that peripheral CD4 T cells from patients with RA were associated with an aberrant STAT3 signaling network and had in fact downregulated *SOCS3* expression (48). We then searched general public repository databases for available transcriptome data from RA synovial tissue. Thirteen studies met our inclusion criteria comprising over 300 synovial samples (**Table S2**). Leveraging all collected publicly available microarray data we specifically looked into the *SOCS3* gene. We found that *SOCS3* was down-regulated in RA synovial tissue compared to controls with the fold change 0.65 and p-value 5E-10 (**Fig. 7A-B)**. Distribution plots per individual study are shown in **Fig. S4**. Furthermore, we interrogated the expression of additional IL-6 signaling pathway components and found minimal to no differences between patients with RA and controls, apart from *JAK2* (**Fig. 7B-C**). These results suggest a mechanism whereby IL-6 signaling may go unchecked and contribute to persistent inflammatory responses in the RA joint.

**Figure 7.**
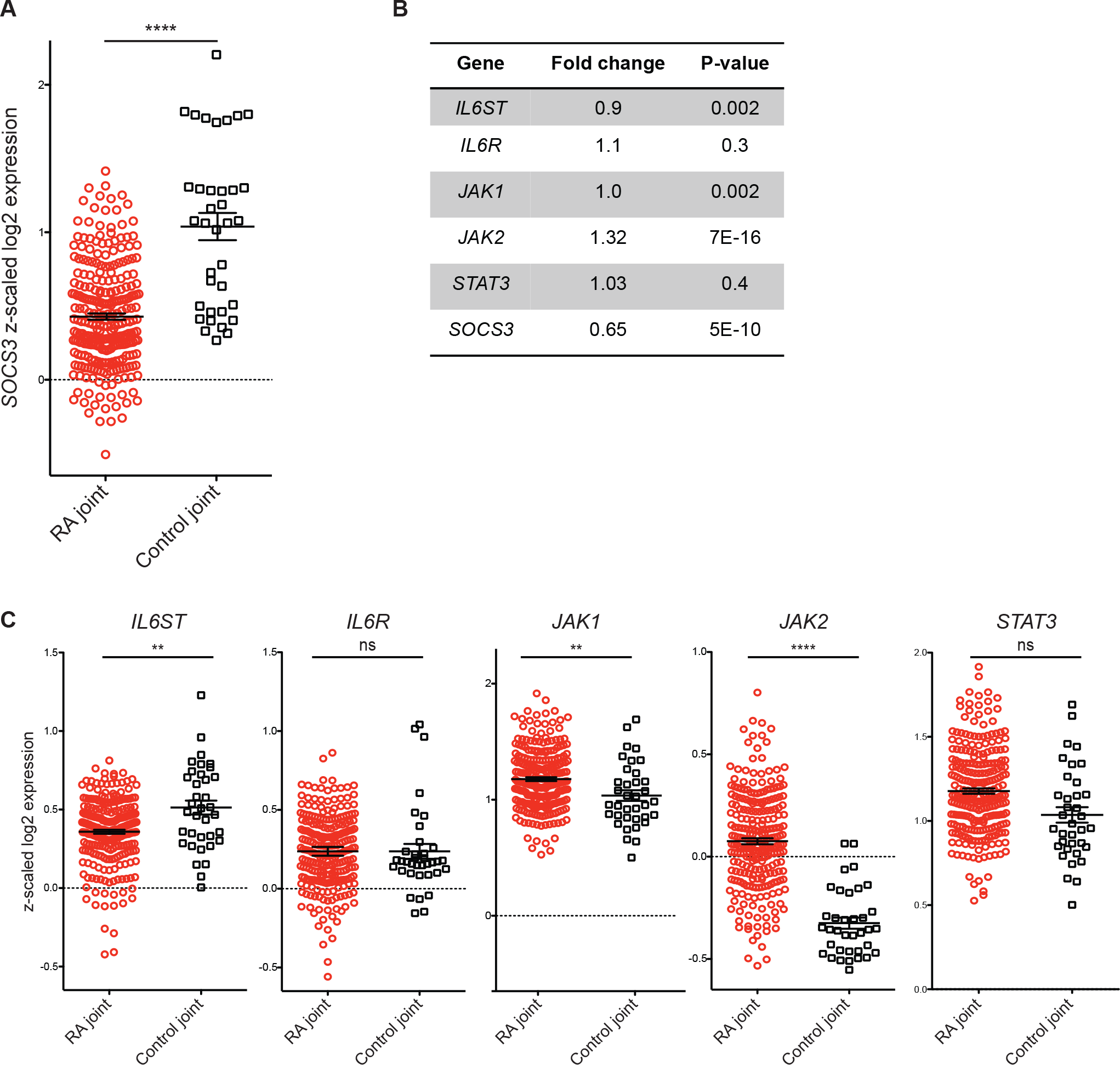
Comparison of gene expression levels in synovium from patients with RA. The gene expression values were log2-transformed and z-scaled. The gene expression levels are summarized in the form of mean ± SEM. **(A)** Synovial *SOCS3* expression levels. **(B)** Table with summary statistics for genes in IL-6 signaling pathway. **(C)** Expression levels for *IL6ST*, *IL6R*, *JAK1*, *JAK2*, and *STAT3* genes. *P*-values were computed by performing the non-parametric two-sample Mann-Whitney-Wilcoxon test. ns: *P* > 0.05, \*\**P* < 0.01, \*\*\*\**P* < 0.0001.

## Discussion

Here we describe an innovative approach, through the detection and monitoring of Nur77 expression, to distinguish putative autoreactive T cells with increased arthritogenic potential from T cells activated by cytokine signaling. Indeed, we find an enrichment of intracellular Nur77^+^ CD4 T cells in both human RA and murine autoimmune arthritis. Although RA synovial tissue has recently been phenotyped with attention placed on infiltrating ‘peripheral helper’ T (Tph) cells, little data exists on infiltrating T cell antigen-reactivity or specificity (49-54). The ability to identify Nur77^+^ CD4 T cells from RA synovial tissue obtained from patients with well established and, in most cases advanced ‘end-stage’ joint disease, suggests that antigen-specific CD4 T cell responses continue to persist in diseased joints (and may challenge our notion of ‘burned-out’ RA). While we were able to exclude Treg cells among the arthritogenic Nur77-GFP high cells in the joints of SKG mice, we could not do so when endogenous Nur77 staining was used in patients. A subset of these synovial Nur77^+^ CD4 T cells may include regulatory T cells, although Tregs make up only ~10% of RA joint infiltrating CD4 T cells (55). Our future studies will further characterize the synovial CD4 T cell subsets that express high levels of Nur77 (*NR4A1*) using single cell RNA sequencing. For clinical investigators, these results highlight the importance of studying the relevant tissues, rather than blood samples, to elucidate antigen-driven events in relevant target tissues.

The ability to use Nur77 to track antigen-activated T cells influenced our decision to take advantage of the Nur77-eGFP reporter to explore functional heterogeneity of the CD4 T cell repertoire before and after frank autoimmune disease onset in the SKG mouse model of arthritis. The SKGNur reporter revealed: 1) SKG thymocytes are able to upregulate a reporter of TCR signaling during thymic positive selection despite their severely impaired TCR signaling capability, presumably due to high affinity self-pMHC TCRs being selected; 2) further TCR stimulation occurs in the periphery (due to chronic auto-antigen stimulation, further repertoire pruning, or both); and, 3) highly autoreactive, arthritogenic T cells, despite (or rather because of) profound deficiency in their TCR signal transduction capacity, are hyper-responsive to IL-6 stimulation, at least in part due to upregulation of IL-6 signaling machinery and decreased SOCS3 expression. This provides a mechanism whereby an autoreactive repertoire with impaired TCR signaling, as seen in the SKG model, is able to break tolerance, differentiate more readily into pathologic Th17 effector cells in the presence of IL-6 and cause arthritis.

That SKG T cells with severely impaired signaling can effectively differentiate into Th17 effector cells that mediate an inflammatory, destructive arthritis akin to RA seems counter-intuitive. Yet, there are multiple reports linking immune deficiency with immune dysregulation and autoimmunity (6, 26, 29, 30, 39, 56-58). It has been suggested that the partial reduction in the number or function of T cells can disturb a ‘tolerogenic balance’ and thereby generate a combination of immunodeficiency and hyper-immune dysregulation (58). The precise mechanism remains elusive, though murine models suggest multiple factors are at play, including selection of an autoimmune TCR repertoire, lymphopenia-induced exaggerated homeostatic proliferation precipitating autoimmune disease, deficient Treg function, resistance to Treg suppression, and T cell dysregulation causing failed induction of anergy in autoreactive T cells despite chronic antigen stimulation (39, 56). In addition to the SKG mouse model, the *Zap70* allelic series and LAT mutated mice also reveal a causal link between impaired TCR signaling associated with immune dysregulation and autoimmunity (35, 59-62). Similarly, mouse models with cytokine signaling defects such as in STAT5A/5B-deficient mice can also serve as examples of immunodeficiency and immune dysregulation and suggest a synergistic role for cytokine signaling in the activation of self-reactive T cells (58, 63).

The link between immunodeficiency and autoimmunity has long been appreciated in humans. Perhaps it is most evident in patients with primary immunodeficiency where up to 20-25% may develop autoimmune cytopenias, RA, or other autoimmune diseases (64-66). The greatest risk has been associated with partial T cell immunodeficiences and common variable immunodeficiency (66). The reverse association has also been described. In the setting of primary autoimmunity, despite presence of activated immune cells mediating disease, observational studies have found an increased susceptibility to serious infections (independent of immune suppressive medications) and premature aging of the immune system, suggesting that impaired immune function might be a primary defect that in turn subverts tolerance (67-71). For example, RA CD4 T cells have the paradoxical ability to differentiate into pathogenic effector cells despite their impaired response to TCR engagement (6, 7, 11-18). Importantly, this paradox is observed in the SKG model of arthritis, making it one of the few mouse models to capture the contribution of an autoreactive TCR repertoire with impaired TCR signal transduction to the pathogenesis of arthritis.

Notably, our results uncover an unexpected mechanism for initiation of autoimmune arthritis involving crosstalk in peripheral T cells between signaling via the TCR and responsiveness to cytokine-dependent cues through the IL-6 receptor. We show not only that chronic TCR stimulation of GFP^hi^ T cells results in compensatory changes in TCR signal transduction and inhibitory co-receptor expression, but also heightened sensitivity to IL-6 that is due at least in part to increased expression of IL-6 signaling machinery, as well as decreased expression of SOCS3. Since IL-6 signaling drives the differentiation of CD4 T cells into arthritogenic Th17 effector cells and its presence is indispensable for arthritis development in SKG mice, our observations provide a mechanistic link between T cell self-reactivity and pathogenicity (36). While IL-6 is critically important for the differentiation of CD4 T cells into IL-17-secreting CD4 helper T cells, it is also possible that IL-6 receptor signaling integrates with TCR signaling to promote survival of autoreactive T cells in the SKG mouse. Limited literature exists on the integration of TCR and IL-6 cytokine signaling in this capacity. Their coincident responses may create a feed-forward loop whereby the presence of IL-6 protects CD4 T cells from activation-induced cell death, promotes their expansion and survival in the periphery and participates in an IL-17A/IL-6 positive-feedback loop promoting autoimmunity (72-74). Further efforts delineating the networks involved in TCR and cytokine signaling integration in the setting of chronic (auto)-antigen-stimulation are underway.

SOCS3 may provide an additional link between IL-6 and TCR signaling networks (75, 76). SOCS3 is the most abundant of the SOCS family member expressed in unstimulated naïve T cells. Its overexpression appears to inhibit TCR signaling and T helper cell proliferation (75, 77), whereas decreased SOCS3 activity has been implicated in murine autoimmune arthritis (81-83) and in human RA (48, 78-80). Furthermore, the suppression of SOCS3 by antigen stimulation of naïve CD4 T cells has been reported following acute TCR stimulation (75, 77). In this study we propose that auto-antigen-dependent stimulation of the skewed TCR repertoire in SKG mice may lead to the observed SOCS3 suppression in their CD4 T cells. Suppression of SOCS3, possibly driven by chronic autoantigen-dependent TCR-signaling, may also result in altered IL-6 and TCR regulation, further promoting the arthritogenic potential of GFP^hi^ cells in the SKGNur mice. Additionally, we identified reports of gene expression in RA synovium and found a highly significant reduction of *SOCS3*, suggesting the aforementioned mechanism may be relevant in RA and related autoimmune diseases. These results will need to be validated in CD4 synovial T cells from patients with RA. Further work delineating the regulation of SOC3 in the setting of chronic (auto)-antigen-stimulation remains to be determined.

Thus, our study highlights the functional importance of identifying and studying the arthritogenic T cell population in SKG mice both before and after frank disease initiation, with translatable implications for human RA. Further dissection of the altered TCR and cytokine signaling networks in arthritogenic human CD4 T cells may expand our understanding of RA pathogenesis, and, in so doing, may reveal targets for novel therapeutic approaches to RA. In addition, our study lays the groundwork for the identification of arthritogenic T cell clones and their receptors as well as bona fide autoantigens not only in the SKG model but also in human RA. Our work also provides a platform for investigators to identify autoantigen-specific T cells in other immunologically-mediated human diseases where the inciting antigen is not known— autoimmunity, cancer and checkpoint-blockade, as well as transplant rejection.

## Methods

### Data Reporting

No statistical methods were used to predetermine sample size. The human studies were not randomized and the investigators were not blinded to allocation during experiments and outcome assessment.

### Human research

Research involving human subjects was performed according to the Institutional Review Board at the University of California, San Francisco through an approved protocol titled Identifying arthritogenic T cells in RA Synovial Tissue Study (IRB #13-11485) with appropriate informed consent as required. Patients with seropositive RA fulfilled the ACR 2010 RA classification criteria.

### Patient Identification for synovial tissue

Through use of the UCSF Clinical Data Research Consultations Office, patients with RA referred to UCSF Orthopedic Surgery and scheduled for joint replacement, synovial biopsy or synovectomy were identified through data extraction of UCSF Medical Center’s electronic medical records and recruited to the study. *Inclusion criteria*: adult patients with seropositive (RF and/or anti-CCP antibody) RA diagnosis undergoing a joint procedure or adult patients undergoing one of the above listed procedures for OA, trauma, or other non-inflammatory condition as part of the control group. No patients were excluded based on gender, age, race, or ethnicity. *Exclusion Criteria:* patients with concomitant joint infection, or diagnosis of another inflammatory arthritis. Clinical variables were abstracted from the records. All records are stored in an encrypted, secure desktop accessible only by HIPAA trained personnel.

### Human PBMC preparation

Whole human blood was collected from patients with RA undergoing joint surgery in BD Vacutainer tubes lined with sodium heparin and gently mixed. Monocytes were isolated by density gradient Ficoll separation as previously described (21).

### Human synovial tissue disaggregation

Synovial tissue was collected from RA and OA patients enrolled and consented under the IRB approved protocol or discarded surgical specimens (OA synovial tissue) from surgical joint procedures. The investigators confirmed the sample was synovial tissue by gross inspection. For single cell suspensions, the synovial tissue was minced with scissors to ~2mm^3^ pieces, which were then digested with Liberase TL (50μg/mL, Roche) and DNAse l (200U/mL, Roche), dissociated using the octoMACS dissociation program (Spin 1000rpm 36 seconds) and then placed at 37°C for 30 minutes rotating at 20 rpm followed by octoMACS dissociation program. The enzymatic reaction was quenched by 10% fetal bovine serum in RPMI (Life Technologies) and debris filtered out using two 70μm strainers. The filtration steps removed large pieces of debris, as well as poorly disaggregated cell clumps. Cells were washed in RMPI complete medium and then counted on a cell counter as above and assessed for viability, followed by FACS staining or cryopreservation.

### Mice

BALB/c and SCID BALB/c mice were purchased from Jackson laboratory, and SKG and Nur77-eGFP tg mice were kindly provided by Shimon Sakaguchi (Kyoto University, Kyoto, Japan) and Julie Zikherman (University of California, San Francisco) respectively. Nur77-eGFP mice were backcrossed to BALB/c more than eight times and crossed to SKG mice to make SKGNur mice. All mice were housed and bred in specific pathogen–free conditions in the Animal Barrier Facility at UCSF according to the University Animal Care Committee and NIH guidelines. All animal experiments were approved by the UCSF Institutional Animal Care and Use Committee.

### *In vivo* treatments

Zymosan A (Sigma-Aldrich) suspended in saline at 10 mg/mL was kept in boiling water for 10 min. Zymosan A solution 2 mg or saline was intraperitoneally injected into 8-12 week-old mice. Adoptive transfer experiments were performed as described (26, 37). The highest and lowest 5% Nur77-eGFP purified CD4 T cells sorted on naïve (CD62L^hi^CD44^lo^) or memory (CD62L^lo^CD44^hi^) markers and depleted of Treg marker CD25 were adoptively transferred by tail vein injection into SCID recipients. To prevent complications of weight loss and early onset of inflammatory bowel disease after adoptive transfer of naïve SKG T cells, WT Tregs were added back in a ratio of 1:3. This attenuated the IBD phenotype prior to the onset of arthritis.

### Histology and histopathology scoring criteria

Ankle joints were fixed in buffered 10% formalin, and paraffin-embedded sections were decalcified and stained with hematoxylin and eosin by the UCSF mouse pathology core. Ankle joints were scored in a blinded fashion using a modified protocol (81). Arthritic paws and ankles were given scores of 0-4 for inflammation and pannus according to the following criteria. Severity of arthritic changes, as well as number of individual joints affected, were considered. Inflammation: 0=normal, 1=minimal synovial and periarticular infiltrate of inflammatory cells, 2=mild infiltration, 3=moderate infiltration with moderate edema, 4=marked-severe infiltration affecting most of the area with marked edema. Pannus: 0=normal, 1=minimal infiltration of pannus in cartilage and subchondral bone, 2=mild infiltration of marginal zone with minor cortical and medullary bone destruction in affected joints, 3=moderate infiltration with moderate hard tissue destruction, 4=marked-severe infiltration with marked destruction of joint architecture, affecting most joints. For each animal, the inflammation and pannus scores were determined for the 4 paws/ankles submitted. A sum total (all joints) animal score was determined. Parameters for the various groups were then compared using Student’s t test with significance at *P* < 0.05.

### Murine synovial tissue preparation

Synovial tissues from ankle joints were digested with 1 mg/mL Collagenase IV (Worthington) in RPMI 1640 medium for 1 h at 37°C; digested cells were filtrated through a nylon mesh to prepare single cell suspensions.

### cDNA Synthesis and qRT-PCR

Total RNA was purified using Trizol (Invitrogen) or the RNeasy Micro kit (QIAGEN). cDNA was synthesized using Superscript III First-Strand Synthesis System (Invitrogen) according to the manufacturer’s instructions. TaqMan Real-Time PCR was performed using TaqMan Real-Time PCR Master Mix (Invitrogen) and TaqMan primer (primer 1: 5′-GCCAATTTCCTTGTCTCCTTG-3′, primer 2: 5′-CTGCTCGAAGATTACGAAATGC-3′),and probe: 5′-/56-FAM/ACTGCCTCT/ZAN/GCTTCCTGCTTTGA/31ABkFQ/-3′ purchased from IDT. An ABI Quantstudio machine (Applied Biosystems) was used for all qPCR assays. The relative abundance of all target genes was normalized to the abundance of an internal reference gene (GAPDH for Nr4a1, and β2 microglobulin for Jak 1 and 2). The following primer sequences were used: Nr4a1 (Nur77) forward, 5′-GGCATGGTGAAGGAAGTTGT-3′, reverse, 5′-TGAGGGAAGTGAGAAGATTGGT-3′; Jak1 primer 7: 5'-GGCTTTCTTAGTGGCTACGAG-3', primer 2: 5'-GAACCACCTCAAGAAGCAGAT-3'; Jak2 primer 7: 5'-AGATCCTTTCGGTTTCACTCAT-3', primer 2: 5'-CAAGTGGAGAGTATGTTGCAGA-3'.

### Antibodies and other reagents

The following fluorescent conjugated antibodies were used: antibodies to human CD3ε-BUV395 or PeCy7 (UCHT1; eBioscience and Biolegend), CD4-PerCPCy5.5 (RPA-T4), CD45-FITC (BD Biosciences), CD45RO-Pacific Blue (UCHL1) (Biolegend) for fluorescence-activated cell sorting (FACS) staining. Mouse antibodies: CD4 PerCp-Cy5.5 (GK1.5), TCRβ PerCp-Cy5.5 (H57-597), and CD62L APC (MEL-14) (TONBO); CD44 FITC (IM7), PD-1 PE-Cy7 (29F.1A12), CD69 PE-Cy7 (1D3), CD5 PerCp-Cy5.5 (53-7.3), CD8α Pacific Blue (53-6.7), TCRβ Pacific Blue (H57-597), CD126 APC (D7715A7), CD130 PE (4H1B35), and TIGIT APC (1GP) (BioLegend); CD44 PE-Cy7 (IM7), CD8α APC-eFluor 780 (53-6.7), CD4 BUV395 (GK1.5), CD44 v450 (IM7), CD62L BV711 (MEL-14), CD73 PE, Ki67 APC (B56) (BD Biosciences); anti-Nur77 (clone 12.14) purified or conjugated to PE (eBiosciences), anti-STAT3 (pY705; clone 4/P) conjugated to PE, and anti-STAT3 (total) PE (BD Biosciences), and anti-IL17A (TC11-18H10.1; Biolegend) for intracellular staining. Human Fc blocking reagent (Miltenyi Biotec) and 2.4G2 mouse blocking reagent were used in all human and mouse stains respectively.

### Flow cytometry and cell sorting

Cells were stained with antibodies of the indicated specificities and analyzed on a BD LSR Fortessa flow cytometer. Cells were sorted to >95% purity using a MoFlo XDP (Beckman Coulter). Flow cytometry plots and analyses were performed using FlowJo v9.9.3 (Tree Star).

### AMLR

10% highest and lowest Nur77-eGFP sorted CD4^+^CD25^-^CD62L^hi^CD44^lo^ naïve T cells from SKGNur and WTNur LNs and spleen were labeled with 5 μM Cell Trace Violet according to the manufacturer’s instructions (Life Technologies) before culturing in a 1:10 ratio with 5×10^5^ BALB/c splenic APCs (prepared via Thy1.2^+^ cell depletion by MACS, Miltenyi Biotec) in a 96-well round-bottom plate in complete RPMI medium for 5 days. Cells were harvested and stained with fixable Live/Dead stain, anti-CD4 and anti-TCR-β and analyzed by flow cytometry. FlowJo Proliferation Platform was used to determine the % Divided.

### Intracellular staining

For Nur77 staining in murine and human cells, assays were performed as previously described (21). For intracellular cytokine staining, LN or spleen cells were stimulated with 50 ng/ml PMA and 1 μM ionomycin in the presence of Golgi-Stop (BD Biosciences) for 5 h, and then stained with anti-CD4 and anti–TCR-β and fixed and permeabilized using BD Cytofix/Cytoperm (BD Biosciences), followed by anti–IL-17 staining. Phospho-Erk and phospho-S6 assays: negatively selected peripheral T cells were rested in serum-free medium for 2 h at 37°C, after which they were stimulated with 5 µg/ml of anti-CD3 (clone 2C11), followed by the addition of 50 µg/ml goat anti-Armenian hamster IgG (Jackson Immunoresearch Laboratory) at 37°C for 2, 5, or 10 min. The reaction was terminated with 2% paraformaldehyde. Cells were pelleted, washed, and stained for surface markers, fixed and permeabilized with 90% ice-cold methanol overnight at −20°C. After permeabilization, Cells from each sample were divided and stained in parallel with anti–phospho-p44/42 MAPK (Thr202/Tyr202) or anti-phospho-S6 (Ser240/244; Cell Signaling Technology), followed by staining with donkey anti–rabbit Ig-APC and anti–CD4-BUV395.

Phospho-STAT staining: cells were stained for surface markers and permeabilized with 90% ice-cold methanol as above. After permeabilization, cells were stained with the indicated phospho-STAT antibody.

### Calcium Flux

Peripheral T cells were incubated with the calcium sensitive dye Indo-1 (Invitrogen) for 30 min at 37°C in DMEM with 2% fetal bovine serum and 25 mM Hepes, washed, and stained with anti-CD4-APC-eFluor780, CD44-PeCy7, CD62L-APC. SKGNur cells were barcoded with CD45-PE. After surface staining, SKGNur and WTNur cells were combined and resuspended 1:1 in DMEM medium and warmed to 37°C for 5 min before stimulation. Baseline Ca^2+^ levels were measured for 30 s before and after the addition of anti-CD3ε mAb (clone 2C11; 1 μg/mL) and for 3 min after the addition of cross-linking antibody goat anti-Armenian Hamster IgG (50 μg/mL; Jackson Immunoresearch Laboratory), followed by ionomycin (1 μM). Ca^2+^ increase was measured using the 405/485 nm emission ratio for Indo-1 fluorescence and displayed as a function of time.

### In vitro proliferation assay

10% highest and lowest Nur77-eGFP sorted CD4^+^CD25^-^ CD62L^hi^CD44^lo^ naïve T cells from SKGNur and WTNur LNs and spleen were labeled with 5 μM Cell Trace Violet according to the manufacturer’s instructions (Life Technologies) before culturing with plate-bound αCD3 at the indicated concentration and 2 ug/mL αCD28 in a 96-well flat-bottom plate in complete RPMI medium for 3 days. Cells were harvested and stained with fixable Live/Dead stain and analyzed by flow cytometry.

### Immunoblot analysis

SKG and WT total CD4 cells were purified from single cell suspensions of spleen and lymph nodes by negative selection with biotinylated antibodies and magnetic beads as previously described (82). Cells were lysed by directly adding 10 % NP-40 lysis buffer to the final concentration of 1% NP40 (containing inhibitors of 2 mM NaVO_4_, 10 mM NaF, 5 mM EDTA, 2 mM PMSF, 10 μg/ml Aprotinin, 1 μg/ml Pepstatin and 1 μg/ml Leupeptin). Lysates were placed on ice and centrifuged at 13,000 *g* to pellet cell debris. Supernatants were run on 10% Bis-Tris Protein Gels (ThermoFisher Scientific) and transferred to PVDF membranes using a Trans-Blot Turb Transfer System (Bio Rad). Membranes were blocked using TBS-T buffer containing 3% BSA, and probed with primary antibodies as described, overnight at 4 °C or 2 h at room temperature. The following day, blots were rinsed and incubated with HRP-conjugated secondary antibodies. Blots were detected using a chemiluminescent substrate and a BioRad Chemi-Doc imaging system.

### Statistics

Results from independent experiments were pooled whenever possible, and all data were analyzed using Prism version 5.0f for Mac (GraphPad Software, La Jolla, California). All data were analyzed by comparison of means using paired or unpaired two-tailed Student's *t* tests. Data in all Fig.s represent mean ± S.E.M. unless otherwise indicated. Differences were considered significant at *P* < 0.05.

### Genomic data collection and preprocessing

A comprehensive search for publicly available microarray data at NCBI Gene Expression Omnibus (83) (GEO) database (http://www.ncbi.nlm.nih.gov/geo/) was performed. The keywords “rheumatoid arthritis”, “synovium”, “synovial”, and “biopsy”, among organism “Homo Sapiens” and study type “Expression profiling by array” were used. The criteria for the dataset exclusion from the search results were the lack of annotations and low probe platforms. We were able to collect 13 datasets with 312 synovial biopsies (**Table S2**). Among them there were 276 samples with RA diagnosis confirmed in later follow-ups and 36 healthy tissue biopsies obtained from joint or trauma surgery. The gene expression measurements were made on 4 platforms from the manufacturers Affymetrix and Illumina, and 2 custom non-commercial chips.

The raw data in the .cel, .txt, and .gpr file formats were downloaded and pre-processing steps were performed using R language (84) version 3.4.4 and the Bioconductor (85) packages *affy* (86) and *limma* (87). They included background correction, log2-transformation, and intra-study quantile normalization. Finally, we performed probe-gene mapping, data merging and cross-study normalization using the ComBat (88) method in R package *sva* (89) in order to minimize the batch effect contribution to the analysis results. After the merge the total number of common genes was 7,078 since some studies were performed on older platforms with lower number of probes. We specifically queried this data for the expression of the *SOCS3* gene. The study GSE3698 did not contain additional genes queried in the IL-6 pathway, therefore this study was removed for analysis of *IL6ST*, *IL6R*, *JAK1*, *JAK2*, *STAT3*.

Code availability: the code is available upon request.

Data availability: The data is available in NCBI GEO repository under assertion numbers GSE12021, GSE21537, GSE24742, GSE36700, GSE3698, GSE39340, GSE45867, GSE48780, GSE55235, GSE55457, GSE55584, GSE57376, GSE77298.

## Author Contributions

J.A. and L.H. were equally involved in designing research studies, conducting experiments, acquiring and analyzing data and preparation of the manuscript. J.A. recruited all study patients, collected, processed, and analyzed human PBMCs and synovial specimens. Y.C., S.Y., and D.C. contributed to conducting experiments and acquiring data. D.R. and M.S. were responsible for the genomic data collection and analysis. L.L. and E.H. contributed synovial tissue specimens. J. Z. was involved in research study design and manuscript preparation. A.W. was involved in research study design, analyzing data, and manuscript preparation.

## Acknowledgements

The authors thank the patients for their invaluable contribution; the UCSF Orthopedic Surgical Department for their collaborative effort in obtaining intra-operative surgical specimens; D. Ludwig from the UCSF Clinical Data Research Consultations Office for patient data extraction; M. Krueger for assistance with patient chart review; S. Sakaguchi and M. Nakamura for providing SKG mice; Z. Wang for cell sorting; A. Roque for animal husbandry.

Funding for this project was provided by the Rheumatology Research Foundation Scientist Development Award and K-Bridge Award (to J.A.); the Rosalind Russell Medical Research Foundation Bechtel Award (to J.A.); the UCSF-Stanford Arthritis Center of Excellence (to J.A.), which is funded in part by the Northern California Arthritis Foundation; the Research Evaluation & Allocation Committee (REAC), School of Medicine, UCSF (to J.A.); UCSF Academic Senate Committee on Research intramural grant (to J.A.); the Arthritis National Research Foundation Award (to J.A.); the National Institutes of Health Awards 2PO1 AI091580 (to A.W.), K08 AR072144 (to J.A.) and K01 AR060807 (to L.-Y.H); and the Rheumatology Research Foundation Within Our Reach grant (to A.W.).

