## Supplemental Figures and Tables for "Reporters of TCR signaling identify arthritogenic T cells in murine and human autoimmune arthritis"

Figure S1

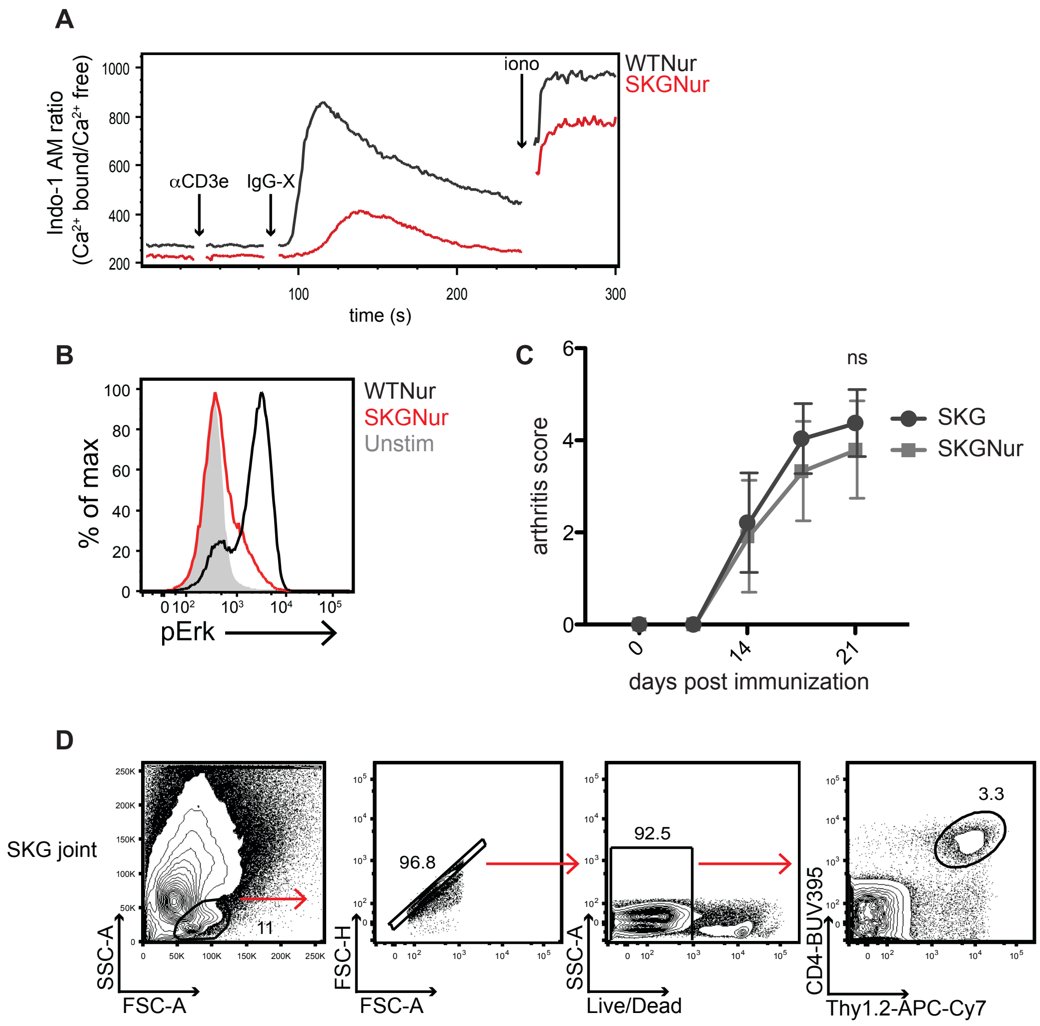


**Fig. S1. SKGNur mice phenocopy SKG mice**

**(A)** TCR-induced calcium increases in CD4$^+$CD25$^-$ T cells from WTNur and SKGNur mice. Data are representative of 4-5 mice in each group from 3 independent experiments.

**(B)** Histograms represent p-Erk levels analyzed in naïve CD4$^+$CD25$^-$ T cells from WTNur and SKGNur mice by flow cytometry after stimulation with α-CD3ε for 2 min. Data are representative of at least 8 mice in each group from 4 independent experiments.

**(C)** Arthritis score in SKG and SKGNur mice after i.p. zymosan immunization, n=7-8 mice in each group.

**(D)** Gating strategy used to identify synovial CD4 T cells in SKG mice by flow cytometry.

Figure S2

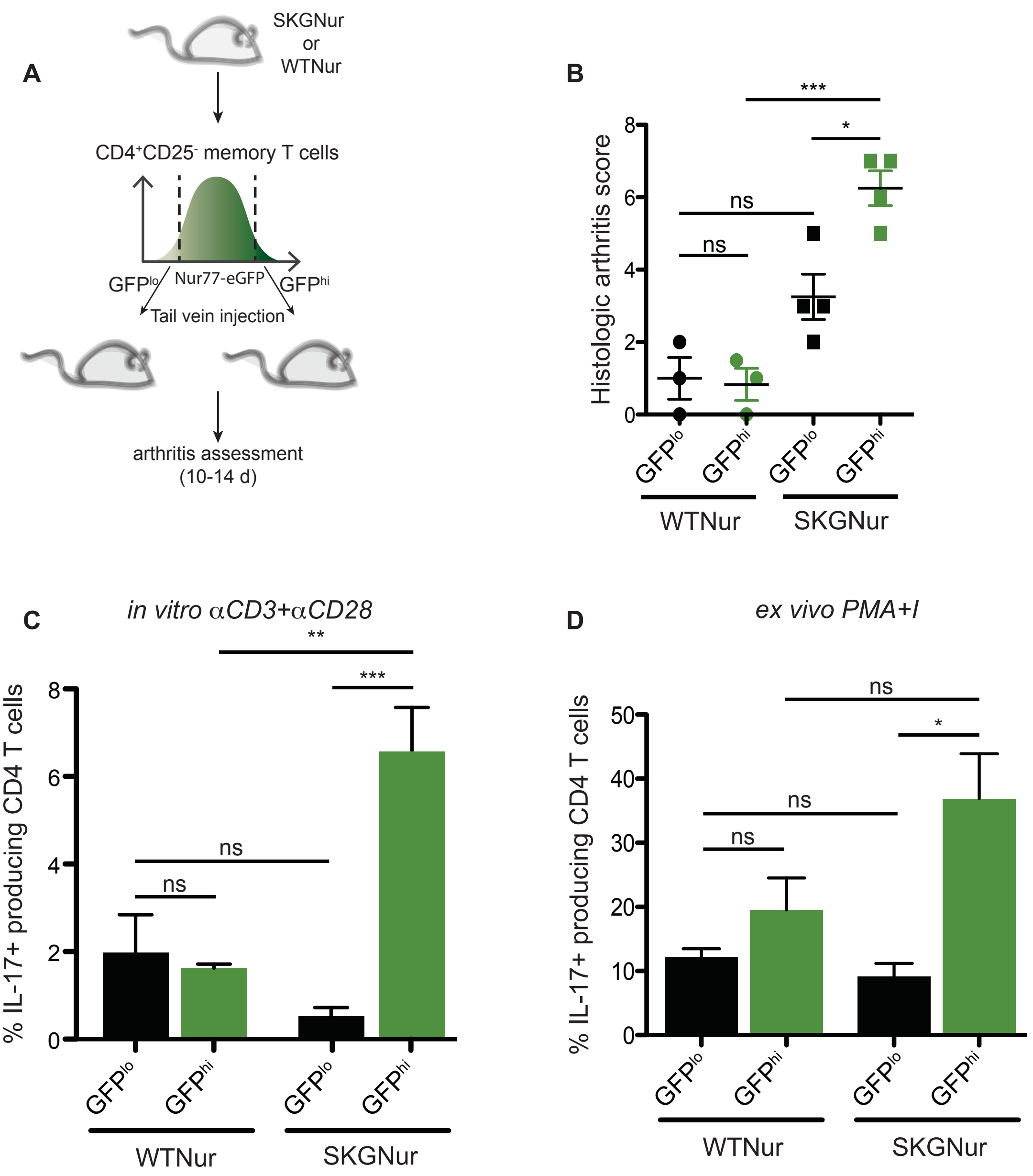


**Fig. S2. Nur77 marks arthritogenicity of CD4 memory T cells in SKGNur mice**
**(A)** The 5% highest and 5% lowest Nur77-eGFP purified CD4 T cells sorted on memory markers (CD62L$^{lo}$CD44$^{hi}$) and depleted of Treg marker CD25 were adoptively transferred into SCID recipients. **(B)** Dot plot of the histology score from SCID recipients that received GFP$^{lo}$ (black) or GFP$^{hi}$ (green) CD4 memory T cells from WTNur (squares) or SKGNur (circles) mice. Each dot represents one biologically independent sample pooled from 2 experiments, presented as mean ± SEM. **(C)** Columns represent sorted WTNur or SKGNur GFP$^{lo}$ (black) or GFP$^{hi}$ (green) naïve CD4 T cells stimulated directly *in vitro* with platebound αCD3 and αCD28 for 6 hours, n=3 per group, mean ± SEM.
**(D)** Columns represent % IL-17 producing GFP$^{+}$ CD4 T cells from SCID recipient splenocytes stimulated i*n vitro* with PMA and ionomycin for 5 hours, mean ± SEM. **(B-D)** ns: $P > 0.05$, *$P < 0.05$, **$P < 0.01$, ***$P < 0.001$ by two-tailed Student's t test.

Figure S3

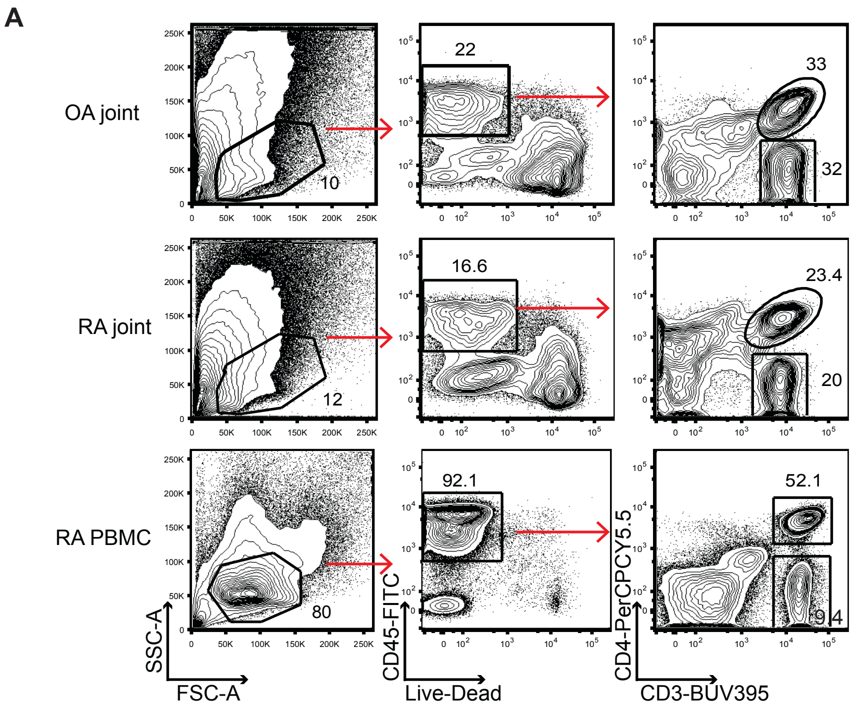


**Fig. S3. Detection of human synovial T cells by flow cytometry**
**(A)** Gating strategy used to identify human CD4 and CD8 synovial and peripheral T cells by flow cytometry.

Figure S4

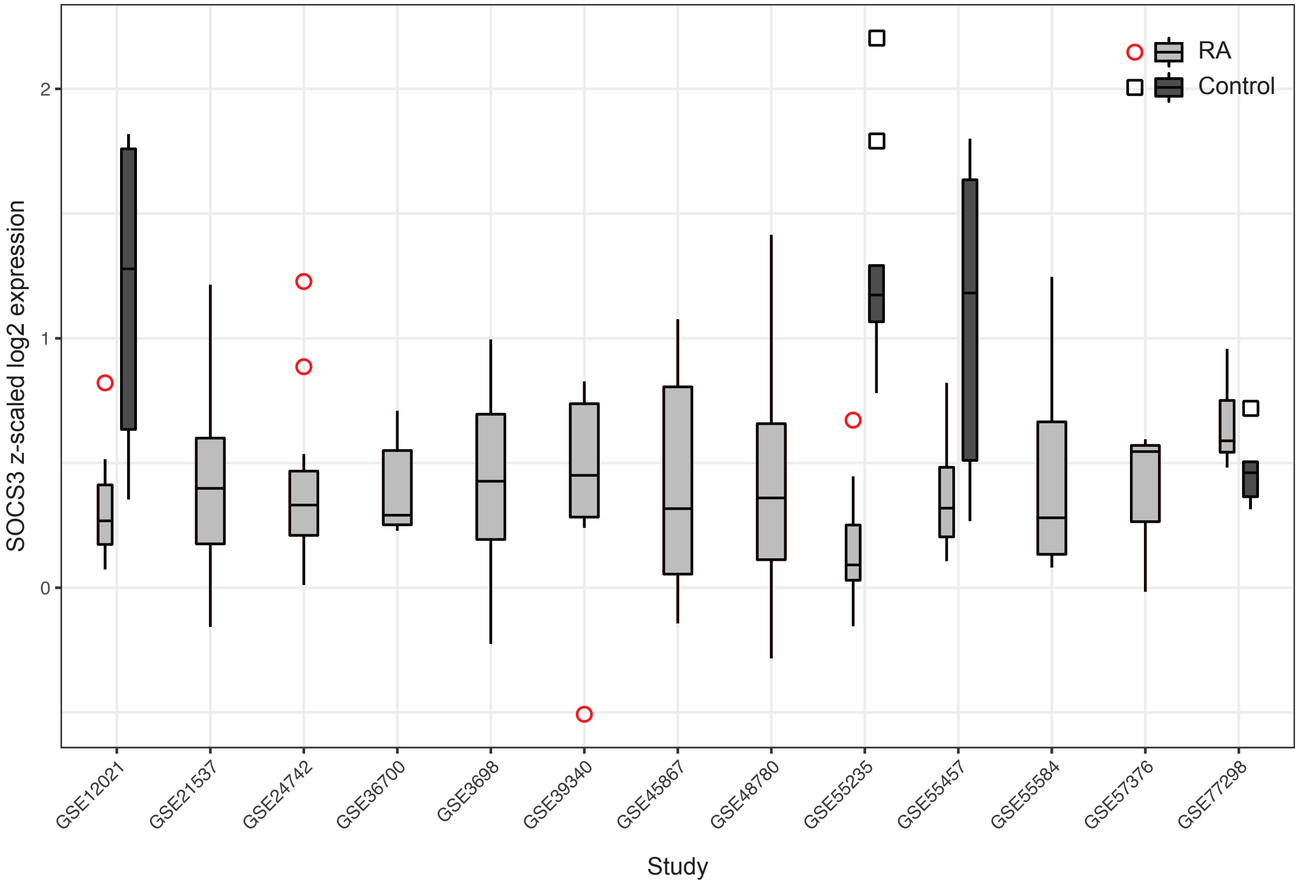


**Fig. S4. SOCS3 expression levels in RA and control synovium by individual study.**
Comparison of *SOCS3* expression levels in synovium from patients with RA and controls per individual study publicly available in NCBI GEO. The expression values were log2 transformed and z-scaled. Boxplots describe the median and inter-quartile range (IQR), whiskers extend to the largest value, but no further than 1.5*IQR, data points beyond whiskers are outlying points.

Table S1

**Table S1. Clinical features of RA patients**
Table summarizing clinical variables abstracted from the records of paitents with seropositive (RF and/or anti-CCP antibody) RA undergoing a joint procedure at UCSF. No patients were excluded based on gender, race, or ethnicity.

**Table 1. Rheumatoid arthritis patient characteristics**

| ID | sex | age | ethnicity | disease duration (yrs) | auto-Ab profile | Biologic | DMARD | prednisone | procedure |
|---|---|---|---|---|---|---|---|---|---|
| 1 | F | 50 | H | 5 | RF, CCP | N | leflunamide | mod | wrist biopsy |
| 2 | F | 60 | C | 41 | RF, CCP | abatacept | MTX | 0 | MCP arthroplasty |
| 3 | F | 58 | C | 15 | RF | rituximab | MTX | mod | MCP arthroplasty |
| 4 | F | 77 | A | 27 | CCP | N | MTX | low | tendon release |
| 5 | F | 68 | A | 14 | RF | N | N | 0 | shoulder arthroscopy |
| 6 | M | 76 | C | 6 | RF, CCP | etanercept | N | low | tendon repair |
| 7 | F | 42 | AA | 22 | RF, CCP | abatacept | N | mod | Hand boutonniere deformity repair |
| 8 | F | 67 | C | 20 | * | N | N | 0 | ankle fusion |

Ethnicity: H = Hispanic, C = Caucasian, A = Asian, AA = African American; N = none;
prednisone dose: 0 = 0 mg/day, low: < 7.5 mg/day, mod: 7.5 < 15 mg/day, high: ≥ 15 mg/day
* auto-Ab profile unavailable; however, clinical features and exam consistent with seropositive RA

Table S2

**Table S2.** Publicly Available Synovium RA Datasets in NCBI GEO. The star * indicates the dataset that was removed in the analysis of additional genes queried in the IL-6 pathway.

| GEO Accession | Platform | Probes | RA | Healthy | Total | Year | Country |
|---|---|---|---|---|---|---|---|
| GSE12021 | GPL96 [HG-U133A] Affymetrix Human Genome U133A Array | 22,283 | 12 | 9 | 21 | 2008 | Germany |
| GSE21537 | GPL7768 KTH Human 30k v1.0 | 36,864 | 62 | 0 | 62 | 2010 | Sweden |
| GSE24742 | GPL570 [HG-U133_Plus_2] Affymetrix Human Genome U133 Plus 2.0 Array | 54,675 | 12 | 0 | 12 | 2010 | Belgium |
| GSE36700 | GPL570 [HG-U133_Plus_2] Affymetrix Human Genome U133 Plus 2.0 Array | 54,675 | 7 | 0 | 7 | 2012 | Belgium |
| GSE3698* | GPL3050 Human Unigene3.1 cDNA Array 37.5K v1.0 | 17,047 | 18 | 0 | 18 | 2005 | Germany |
| GSE39340 | GPL10558 Illumina HumanHT-12 V4.0 expression beadchip | 47,314 | 10 | 0 | 10 | 2012 | China |
| GSE45867 | GPL570 [HG-U133_Plus_2] Affymetrix Human Genome U133 Plus 2.0 Array | 54,675 | 20 | 0 | 20 | 2013 | Belgium |
| GSE48780 | GPL570 [HG-U133_Plus_2] Affymetrix Human Genome U133 Plus 2.0 Array | 42,739 | 83 | 0 | 83 | 2013 | USA |
| GSE55235 | GPL96 [HG-U133A] Affymetrix Human Genome U133A Array | 22,283 | 10 | 10 | 20 | 2014 | Germany |
| GSE55457 | GPL96 [HG-U133A] Affymetrix Human Genome U133A Array | 22,283 | 13 | 10 | 23 | 2014 | Germany |
| GSE55584 | GPL96 [HG-U133A] Affymetrix Human Genome U133A Array | 22,283 | 10 | 0 | 10 | 2014 | Germany |
| GSE57376 | GPL13158 [HT_HG-U133_Plus_PM] Affymetrix HT HG-U133+ PM Array | 54,715 | 3 | 0 | 3 | 2014 | USA |
| GSE77298 | GPL570 [HG-U133_Plus_2] Affymetrix Human Genome U133 Plus 2.0 Array | 54,675 | 16 | 7 | 23 | 2016 | Netherlands |
| | | **TOTAL:** | 276 | 36 | 312 | | |
